# SORLA-driven endosomal trafficking regulates the oncogenic fitness of HER2

**DOI:** 10.1101/299586

**Authors:** Mika Pietilä, Pranshu Sahgal, Emilia Peuhu, Niklas Jäntti, Ilkka Paatero, Olav M. Andersen, Artur Padzik, Matias Blomqvist, Immi Saarinen, Peter Boström, Pekka Taimen, Johanna Ivaska

## Abstract

Human epidermal growth factor receptor 2 (HER2) is an oncogene targeted by several kinase inhibitors and therapeutic antibodies. Endosomal trafficking of many other receptor tyrosine kinases regulates their oncogenic signaling, but the prevailing view is that HER2 is retained on the cell surface. Here we reveal that in cancer cells Sortilin related receptor 1 (SORLA; *SORL1*) forms a complex with HER2 and regulates its subcellular distribution by promoting recycling of endosomal HER2 back to plasma membrane. Expression of SORLA in cancer cell lines and bladder cancers correlates with HER2 levels. Depletion of SORLA targets HER2 to late endosomal/lysosomal compartments, impairs HER2-driven signaling and *in vivo* tumor growth. SORLA silencing also disrupts normal lysosome function and sensitizes anti-HER2 therapy sensitive and resistant cancer cells to lysosome-targeting cationic amphiphilic drugs. These findings reveal potentially important SORLA-dependent endosomal trafficking-linked vulnerabilities in HER2-driven cancers.

## Introduction

The human epidermal growth factor receptor 2 (HER2; also known as ErbB2) is a receptor tyrosine kinase and a well-established oncogene. HER2 amplification is found in 15-30 % of breast cancers ^1, 2^, and HER2 overexpression or activating mutations are clinically relevant in other solid tumors, such as bladder cancer, and colorectal, gastric and lung adenocarcinoma ^3–5^. The biological relevance of HER2 as a driver oncogene is undisputed, and several targeted therapies have been approved for treating HER2-dependent cancers.

Several previous studies, including ours, demonstrate that oncogenic signaling and endosomal traffic of many receptor tyrosine kinases are functionally coupled ^6–9^. Endosomal trafficking critically controls, for instance, the strength and the duration of signaling by epidermal growth factor receptor (EGFR; also known as ErbB1) or MET ^6, 10, 11^. However, in comparison to other receptor tyrosine kinases, the details of HER2 trafficking are poorly understood. The prevailing view, supported by two different models, is that HER2 resides almost exclusively on the plasma membrane in HER2-amplified cancer cell lines ^12^. Data in favor of a ‘lack of internalization’ model suggest that HER2 resists internalization due to 1) absence of identified ligands required for induction of endocytosis ^12^, 2) absence of a recognizable internalization motif ^12^, 3) inhibition of clathrin-coated pit formation ^13^, 4) association with membrane protrusions ^14^ and 5) heat shock protein 90 (HSP90)-dependent stabilization of HER2 on the plasma membrane ^15–17^. On the other hand, data in support of a model for ‘rapid recycling’ resulting in nearly exclusive cell-surface localization of HER2 have been brought forward ^18–20^. Hence, the key outstanding questions are: does HER2 undergo endosomal trafficking in HER2-driven cancer cells, and what would the functional consequences of HER2 trafficking be for its oncogenic properties?

SORLA is a multifunctional intracellular sorting protein belonging to sortilin and LDL-receptor families consisting of a large extracellular domain, a transmembrane domain and a short cytoplasmic tail ^21^, which is essential for its membrane sorting functions ^22, 23^. SORLA has been implicated in regulating amyloid precursor protein (APP) processing during the pathogenesis of Alzheimer’s disease (AD) ^24–26^ and in lipid metabolism and obesity ^27, 28, 29, 30^. We made the unexpected observation that SORLA is highly expressed specifically in HER2-driven cancers where SORLA levels correlate with HER2 subcellular localization. We find that SORLA associates with HER2, and regulates its trafficking from intracellular compartments to the plasma membrane. Moreover, we show that SORLA is necessary for oncogenic HER2 signaling *in vitro* and *in vivo*. Depletion of SORLA induces lysosome dysfunction and results in mislocalization of signaling-defective HER2 to lysosomes. As a result, SORLA depletion halts tumorigenesis and sensitizes cancer cells to a clinically relevant lysosome-targeting drug. Taken together, our data demonstrate that SORLA has previously unrecognized oncogenic functions in carcinomas.

## Results

### SORLA associates and co-localizes with HER2 on the plasma membrane and intracellular vesicles

Analysis of different breast cancer cell lines revealed that SORLA protein was mainly expressed in those with HER2-amplification (Fig. 1a). SORLA was also highly expressed in the 5637 bladder cancer cell line harboring a HER2-activating mutation (S310F) ^31^ when compared to the HER2-low expressing T24 cell line and primary patient-derived bladder cancer cells (Supplementary Fig. 1a). As SORLA has not previously been scrutinized in carcinoma cells, we first examined the subcellular localization of this sorting receptor in different endosomal compartments. SORLA was found to localize largely to early endosomes (identified by EEA-1- and Rab5-positive expression) and retrograde vesicles (VPS35) in breast cancer cells. SORLA-GFP fusion protein showed similar localization as endogenous SORLA to EEA1- and VPS35-positive vesicles (Supplementary Fig. 1b) and was also present in recycling endosomes (Rab11) (Supplementary Fig. 1b) but did not overlap with late-endosome or lysosome markers Rab7 and LAMP1, respectively (Supplementary Fig. 1b).

**Figure 1.**
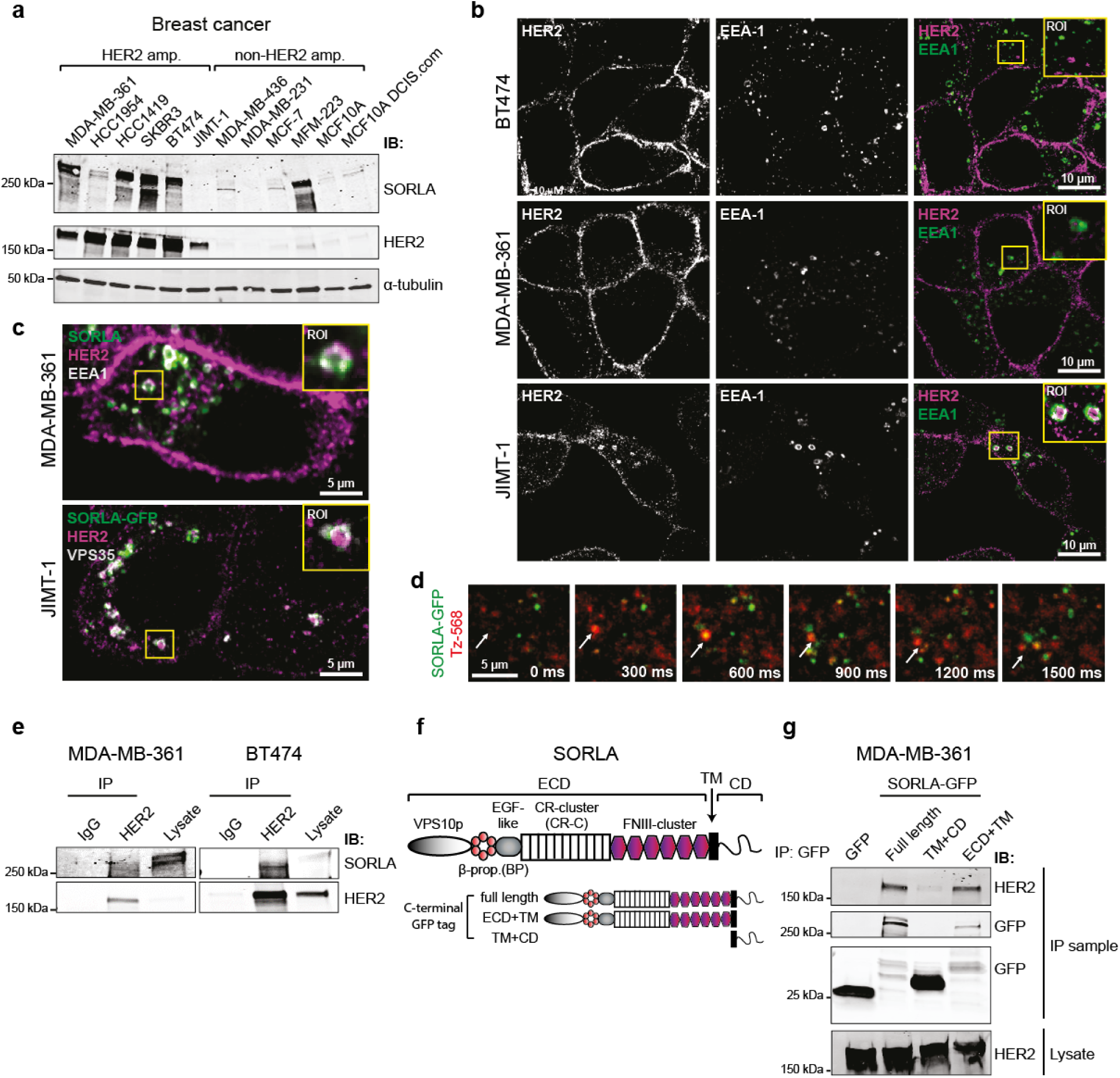
SORLA is highly expressed in HER2 breast cancer cells and associates and co-traffics with HER2 (**a**) Western blot analysis of SORLA and HER2 in breast cancer cell lines. α-tubulin is a loading control. (**b**) Confocal microscopy imaging of HER2 (magenta) and EEA-1 (green) in BT474, MDA-MB-361 and JIMT-1 cells. (**c**) Endogenous SORLA (green), HER2 (magenta) and EEA-1 (white) staining in MDA-MB-361 cells (top panel). Endogenous HER2 (magenta) and VPS35 (white) staining in JIMT-1 cells expressing SORLA-GFP (green) (bottom panel). (**d**) Live-cell TIRF imaging of AlexaFluor-568-labeled trastuzumab (Tz-568; red) and SORLA-GFP (green) in MDA-MB-361 cells. (**e**) Co-immunoprecipitation of endogenous SORLA with endogenous HER2 in MDA-MB-361 and BT474 cells. (**f**) Schematic of the SORLA protein domains and summary of the constructs used. (**g**) Co-immunoprecipitation of endogenous HER2 with different SORLA-GFP fragments in MDA-MB-361 cells. Where immunoblots and micrographs are shown, these are representative of n = 3 independent experiments; IB: immunoblotting, IP: Immunoprecipitation. ECD: extracellular domain; TM: transmembrane domain; CD: cytosolic domain.

Previous studies have indicated that HER2 is mainly restricted to the plasma membrane ^12^, but these observations were based on a very limited number of cell lines and/or on exogenous overexpression of HER2 ^14, 18, 32^. Since SORLA is an important sorting protein in neuronal cells and adipocytes and is expressed in HER2-positive cancer cells (Fig. 1a), we were interested to re-examine the subcellular localization of endogenous HER2 in breast cancer cells with variable levels of SORLA expression (Fig. 1a). In the BT474 and SKBR3 cells with high SORLA expression, HER2 was mainly on the plasma membrane, in accordance with previous work ^14, 18^, and did not overlap with EEA-1 positive endosomes (Fig. 1a,b, Supplementary Fig. 1c). A similar distribution was also detected in the third SORLA-high cell line HCC1419 (Fig. 1a, Supplementary Fig. 1c). However, in cells with intermediate (MDA-MB-361) or low/absent (HCC1954 and JIMT-1) SORLA, HER2 was also distributed intracellularly indicating that HER2 must be trafficking at least in a subset of HER2 cancers. In these cells, a proportion of the intracellular HER2 pool clearly overlapped with EEA-1 indicating localization in early endosomes (Fig. 1b, Supplementary Fig. 1c). In MDA-MB-361 cells endogenous SORLA co-localized with HER2 in EEA-1 positive early endosomes (Fig. 1c), and in the SORLA negative JIMT-1 cells SORLA-GFP localized to HER2 and VPS35 positive intracellular vesicles (Fig. 1c). When SORLA-GFP expression was analyzed in JIMT-1 cells (n = 130), 19.8±0.8 percent of SORLA-GFP overlapped with intracellular HER2 (Supplementary Fig. 1d) showing Pearson’s R of 0.3±0.01 indicative of partial colocalisation.

To study whether SORLA and HER2 would show similar dynamics in cells, we chose to image the MDA-MB-361 cells expressing intracellular as well as cell surface pools of HER2. We performed live-cell TIRF imaging (allowing visualization of events close to the plasma membrane) of SORLA-GFP and HER2 labelled with an Alexa568-conjugated anti-HER2 antibody (trastuzumab; Tz-568). Short-lived SORLA- and HER2-positive structures were detected in the TIRF-plane, indicative of active dynamics to and from the plasma membrane. In addition, co-localizing puncta of SORLA and HER2 were frequently observed undergoing dynamic lateral movement on the plasma membrane (Fig. 1d, Supplementary video 1). Live-cell imaging deeper in the cytoplasm showed that SORLA and HER2 move together within the same endosomal structures (Supplementary Fig. 1e and video 2). Collectively these data show that SORLA and HER2 undergo co-trafficking between the plasma membrane and endosomes.

Intrigued by the apparent co-trafficking of SORLA and HER2, we next performed a set of immunoprecipitation assays to investigate whether HER2 and SORLA associate. Endogenous HER2 co-precipitated endogenous SORLA in MDA-MB-361 and BT474 cells, indicating that HER2 and SORLA are in the same protein complex (Fig. 1e). SORLA consists of an extracellular domain (ECD), a transmembrane domain (TM) and a short cytosolic domain (CD) (Fig. 1f). To study the link between HER2 binding and different domains of SORLA in cells, we generated truncated SORLA-GFP fusions consisting of either the SORLA extracellular and transmembrane domains (ECD+TM) or the SORLA transmembrane and cytosolic domains (TM+CD) (Fig 1f, g). HER2 associated with the full-length SORLA-GFP and SORLA-GFP ECD+TM in cells, but failed to associate with SORLA-GFP TM+CD (Fig. 1g). Interestingly, SORLA-GFP TM+CD showed similar vesicular localization as full-length SORLA-GFP, whereas SORLA-GFP ECD+TM was found diffusely in membrane compartments in the cytoplasm and on the plasma membrane (Supplementary Fig. 1f). Thus, SORLA ECD is necessary for association with HER2, whereas SORLA CD is responsible for correct subcellular localization of the protein.

### SORLA regulates HER2 cell surface levels and HER2 oncogenic signaling

The apparent inverse correlation between SORLA levels and the proportion of intracellular HER2 in the different HER2 cell lines (Fig 1a, b Supplementary Fig. 1c) prompted us to hypothesise that cell surface HER2 levels may be regulated by SORLA. To test this we performed loss-of-function experiments in SORLA-high BT474 cells and gain-of-function experiments in intermediate/negative SORLA cell lines MDA-MB-361 and JIMT-1 cells, respectively. In BT474 cells, with predominantly plasma membrane localized HER2 and high SORLA expression, silencing of SORLA led to, approximately, a 50 % decrease in cell-surface HER2 protein levels (Fig. 2a). Conversely, in the SORLA intermediate MDA-MB-361 and SORLA-negative JIMT-1 cells, in which HER2 localizes more to endosomal structures, SORLA overexpression significantly increased cell surface HER2 levels (Fig. 2a). Total HER2 protein levels followed a similar trend of being significantly downregulated in SORLA silenced BT474 cells and upregulated in SORLA-overexpressing MDA-MB-361 and JIMT-1 cells (Fig. 2b, c). Quantitative PCR analysis of *ErbB2* mRNA levels after SORLA silencing or overexpression did not show any significant difference indicating that SORLA-mediated regulation of HER2 occurs predominantly at the post-transcriptional level (Supplementary Fig. 2a).

**Figure 2.**
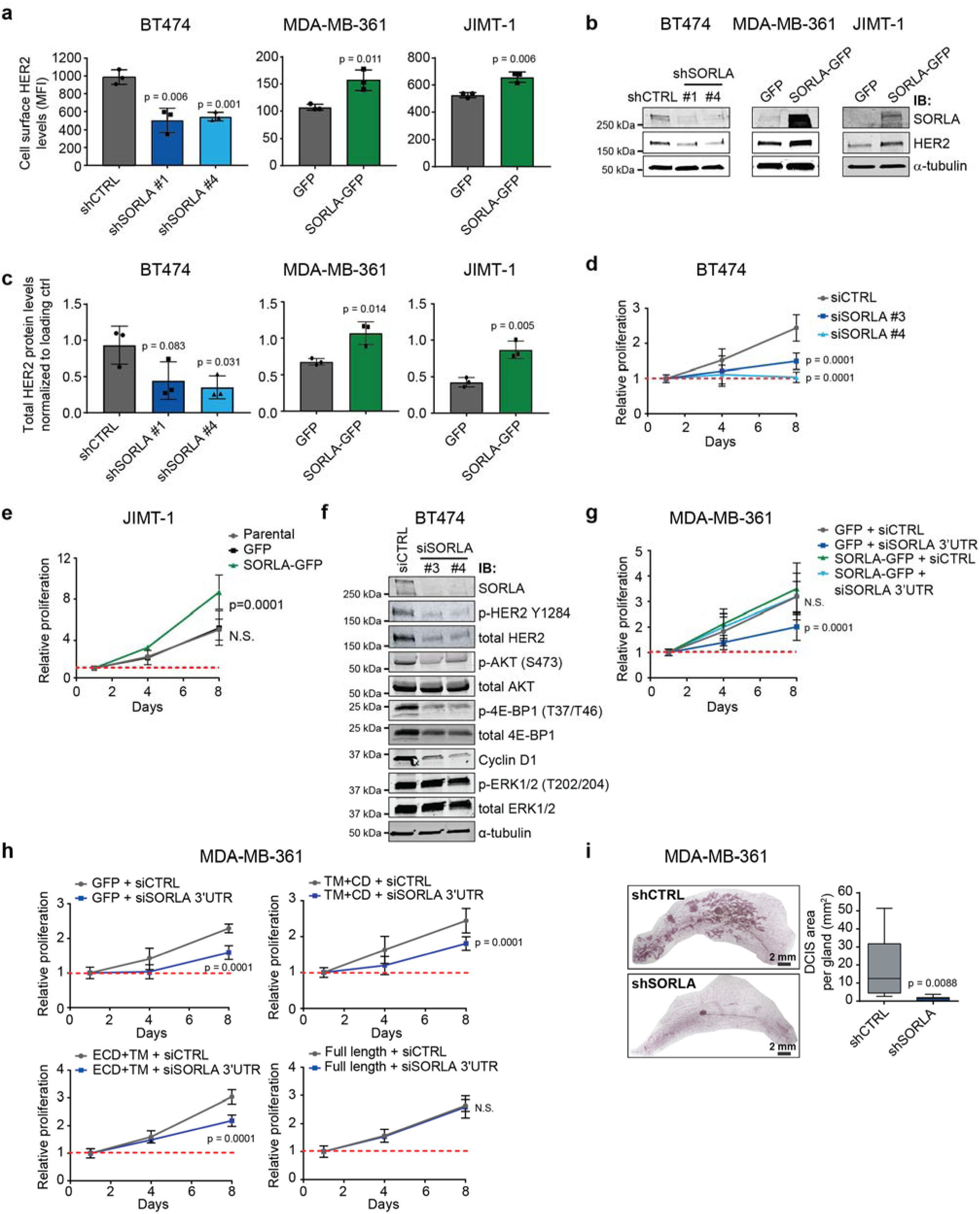
SORLA regulates cell surface HER2 levels and oncogenic signaling in breast cancer cells. (**a**) Flow cytometry analysis of HER2 levels on the cell surface in the indicated cell lines after silencing with control (shCTRL) and SORLA (shSORLA #1 and #4) shRNAs and overexpression of control-GFP and SORLA-GFP; SORLA-high BT474 cells with predominantly plasma membrane HER2 were chosen for the silencing and SORLA low/negative MDA-MB-361 and JIMT-1 cells with endosomal HER2 were selected for the SORLA-GFP overexpression experiments (mean ± s.d of n = 3 independent experiments; statistical analysis: unpaired Student’s *t*-test). (**b**) Immunoblotting of SORLA and HER2 protein levels in the indicated transfectants. α-tubulin is a loading control. (**c**) Quantification of total HER2 protein levels in the indicated transfectants (mean ± s.d of n = 3 independent experiments; statistical analysis: unpaired student’s t-test). (**d**) Proliferation of BT474 cells after SORLA silencing with siRNAs (mean ± s.d of n = 12 (4 technical replicates, 3 independent experiments); statistical analysis: two-way ANOVA). (**e**) Proliferation of parental, GFP control and SORLA-GFP overexpressing JIMT-1 cells (mean ± s.d of n = 12 (4 technical replicates, 3 independent experiments); statistical analysis: two-way ANOVA). (**f**) Western blot analysis of the indicated signaling proteins (phosphorylated and total) in BT474 cells after scramble (siCTRL) and SORLA silencing (siSORLA #3 and siSORLA #4), α-tubulin is a loading control. (**g**) Proliferation of MDA-MB-361 cells expressing SORLA-GFP or GFP control after silencing with control siRNA or siRNA against the 3’UTR of SORLA (mean ± s.d of n = 14 from 4 independent experiments, two-way ANOVA). (**h**) Proliferation of MDA-MB-361 cells expressing SORLA-GFP constructs after silencing with control siRNA (siCTRL) or siRNA against the 3’UTR of SORLA (siSORLA 3’UTR). (mean ± s.d. of n = 8 from two independent experiments; statistical analysis: two-way ANOVA). (**i**) Quantification of mammary DCIS tumors (Carnoy-stainings) 10 weeks after injection of shSORLA and shCTRL silenced MDA-MB-361 cells into mammary ducts of NOD.SCID mice (box plot represents median and 25^th^ and 75^th^ percentiles (interquartile range; IQR) and whiskers extend to maximum and minimum values; n = 9 mice per group; statistical analysis: unpaired Student’s *t*-test). Where immunoblots are shown, these are representative of n = 2-5 independent experiments; IB: immunoblotting.

HER2 amplification is a major driver of proliferation and tumorigenesis, which prompted us to explore whether SORLA plays a functional role in breast cancer cells. Efficient silencing of SORLA in HER2-amplified SORLA-high BT474 cells significantly reduced cell proliferation (Fig. 2d, Supplementary Fig. 2b). This was specifically due to loss of SORLA rather than off-target-effects given that silencing SORLA with five individual siRNAs significantly reduced proliferation in BT474 cells (Supplementary Fig. 2c, d). Conversely, in the SORLA negative JIMT-1 cells expression of SORLA-GFP, which triggered HER2 up-regulation on the plasma membrane (Fig. 2a), significantly increased proliferation of these cells (Fig. 2e, Supplementary Fig. 2e). When SORLA was silenced in non-HER2-amplified FGFR2-amplified MFM-223 breast cancer cells (Fig. 1a), there was no effect on cell proliferation (Supplementary Fig. 2f), indicating that SORLA is required for proliferation only in HER2-dependent cancer cells.

HER2 signaling on the plasma membrane along the PI3K/AKT pathway is critical for HER2 growth promoting functions in cancer cells. Silencing of SORLA in BT474 and MDA-MB-361 cells led to decreased phosphorylation of AKT (Ser473) and 4E-PB1 (Thr37/46) as well as decreased cyclin D1 levels, but did not inhibit mitogen-activated protein kinase (ERK1/2) signaling (Fig. 2f Supplementary Fig. 2g). Taken together these data suggest that SORLA silencing specifically attenuates cell proliferation and PI3K-dependent HER2 signaling in HER2-amplified cells.

To investigate further the requirement for SORLA for proliferation of HER2 cancer cells, we silenced SORLA in MDA-MB-361 cells with intermediate SORLA expression using a SORLA 3’UTR targeting siRNA and then re-expressed SORLA. Re-expression of SORLA-GFP in MDA-MB-361 cells fully rescued cell proliferation to control levels (Fig. 2g, Supplementary Fig. 2h). Importantly, only full-length SORLA-GFP, and not the HER2-associating SORLA-GFP ECD+TM or the correctly localizing but none-HER2-binding SORLA-GFP TM+CD, was sufficient to rescue the effect of SORLA silencing on MDA-MB-361 cell proliferation (Fig. 2h). This suggests that SORLA association with HER2 and SORLA sorting functions are necessary for SORLA-mediated proliferation of HER2-dependent cancer cells.

Importantly, silencing SORLA compromised the *in vivo* tumor engraftment of HER2-amplified breast cancer cells in an orthotropic model. SORLA-silenced and control MDA-MB-361 cells were generated by transducing two short hairpin RNAs (shRNAs) targeting SORLA (shSORLA #1 and shSORLA #4) and a non-targeting control (shCTRL). Efficient SORLA silencing strongly inhibited *in vitro* proliferation of these cells (Supplementary Fig. 2i, j). When control-silenced cells were injected into mammary ducts of immunocompromised NOD.SCID mice, multiple ductal carcinoma in situ (DCIS) lesions were formed within 10 weeks. In contrast, the development of DCIS tumors from SORLA-silenced xenografts was almost completely halted (Fig. 2i). Take together these data indicate that SORLA functionally regulates both the expression and oncogenic functions of HER2 in breast cancer.

### SORLA promotes HER2 recycling

The steady-state distribution of cell surface receptors between endosomes and the plasma membrane is regulated by the respective rates of receptor internalization and recycling back to the plasma membrane. To study whether SORLA regulates this balance in HER2 cell lines, we investigated HER2 localization and trafficking in the presence and absence of SORLA. First, we silenced SORLA in BT474 cells, with high SORLA levels and predominantly plasma membrane HER2 at steady state. Interestingly, shRNA-mediated SORLA silencing led to increased intracellular accumulation of HER2, normally not observed in these cells (Fig. 3a, b). A similar shift in HER2 subcellular localization was triggered when inhibiting vesicular recycling with primaquine, indicating that HER2 undergoes constant endocytosis balanced with very rapid recycling in cells with predominantly plasma membrane localized HER2 and that SORLA may play a role in facilitating this (Fig 3c, d).

**Figure 3.**
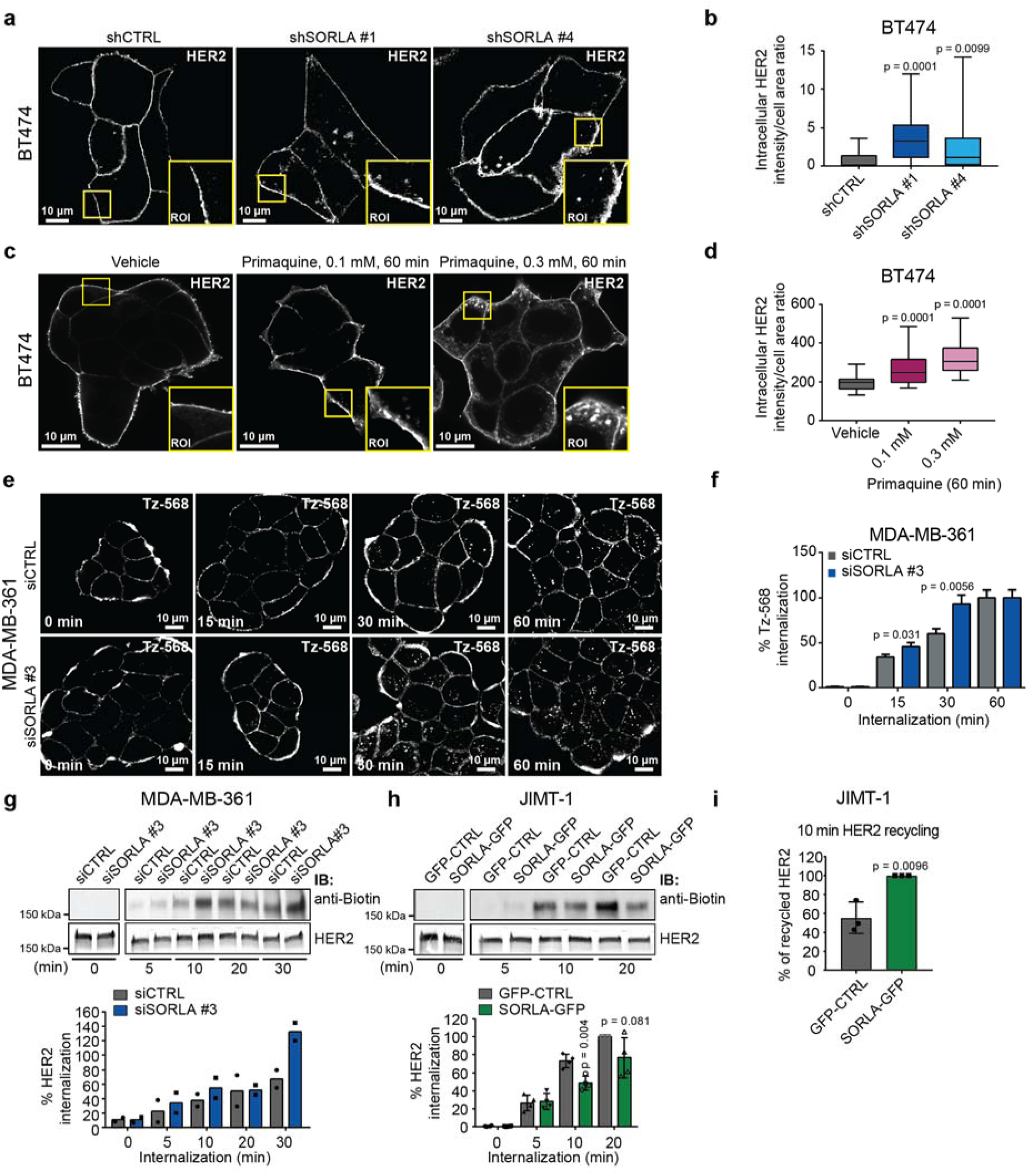
SORLA promotes HER2 recycling. (**a**) Confocal images of HER2 expression after SORLA silencing with shRNAs (shSORLA #1 and shSORLA #4) in BT474 cells. shCTRL: scramble shRNA silenced controls. (**b**) Quantification of intracellular HER2 signal from (a) displayed as a box plot (n = 20-31 cells from two experiments, analysis performed on 8-bit images; statistical analysis: Mann-Whitney test). (**c**) Confocal images of HER2 in vehicle and primaquine treated (for 60 min) BT474 cells. (**d**) Quantification of intracellular HER2 signal from (c) displayed as a box plot (n = 23-35 cells, analysis performed on 16-bit images; statistical analysis: Mann-Whitney test). (**e**) Microscopy analysis of AlexaFluor 568-labeled trastuzumab (Tz-568) internalization after 0 - 60 min in MDA-MB-361 cells silenced with SORLA (siSORLA #3) or scramble (siCTRL) siRNA. (**f**) Quantification of intracellular Tz-568 signal from (e) (mean ± s.e.m; n = 64-111 cells from two independent experiments; statistical analysis: Mann-Whitney test). (**g**) Immunoblotting analyses of biotin-labeled cell-surface HER2 internalization after 0 - 30 min in MDA-MB-361 cells treated with scramble (siCTRL) and SORLA (siSORLA #3) siRNAs, and quantification of the internalized HER2 relative to total HER2 (data are mean ± range of n = 2 independent experiments). (**h**) Immunoblotting analyses of biotin-labeled cell-surface HER2 internalization after 0 - 20 min in JIMT-1 cells overexpressing SORLA-GFP (or control GFP; GFP-CTRL), and quantification of the internalized HER2 relative to total HER2 (data are mean ± s.d.; n = 4 independent experiments; statistical analysis: unpaired Student’s *t*-test). (**i**) Quantification of HER2 recycling rate (return of internalized biotinylated cell-surface HER2 back to the plasma membrane) in JIMT-1 cells transfected with GFP-CTRL or SORLA-GFP (data are mean ± s.d.; n = 3 independent experiments; statistical analysis: unpaired Student’s *t*-test). Box plots represent median and 25^th^ and 75^th^ percentiles (interquartile range; IQR) and whiskers extend to maximum and minimum values. Where immunoblots and micrographs are shown, these are representative of n = 3 independent experiments; IB: immunoblotting; ROI: magnified region of interest.

Next we investigated HER2 dynamics in MDA-MB-361 cells where HER2 is localized endosomally and on the plasma membrane and SORLA is expressed at intermediate levels. SORLA-silenced MDA-MB-361 cells were subjected to an imaging-based receptor uptake assay. Cell surface HER2 was labelled with Tz-568 on ice, after which internalization of Tz-568 was induced by incubating cells at +37°C for 15 min, 30 min, and 60 min before fixing. SORLA-silenced cells showed significantly greater accumulation of intracellular HER2 after 15 min and 30 min of internalization when compared to control-silenced cells (Fig. 3e, f). Similar data was obtained with a biochemical internalization assay of HER2. SORLA-silenced MDA-MB-361 cells were cell surface biotinylated on ice with a cleavable biotin-conjugate. Internalization was induced by incubating cells at +37°C for the indicated times and was followed by cleavage of cell surface retained biotin before cell lysis, HER2 immunoprecipitation and detection of biotinylated and total HER2 with anti-biotin and anti-HER2 antibodies, respectively. In line with the imaging-based internalization analyses, loss of SORLA resulted in increased intracellular accumulation of HER2 at 30 min internalization (Fig. 3g), indicating that the outcome of the imaging-based internalization assays was not secondary to trastuzumab-induced HER2 uptake ^18^. In most systems, receptor endocytosis is most prominent immediately after synchronized internalization and reaches a plateau after 10 minutes in cells where receptor recycling is unperturbed. Since SORLA silencing caused delayed HER2 accumulation inside MDA-MB-361 cells and the recycling inhibitor primaquine triggered intracellular accumulation of HER2 in BT474 cells, we hypothesized that SORLA could regulate the recycling step of HER2 trafficking. To test this in more detail, we utilized JIMT-1 cells which do not express endogenous SORLA, contain a substantial fraction of intracellular HER2 at steady state and where overexpression of SORLA leads to increased HER2 cell surface levels (Fig 2a). Overexpression of SORLA-GFP in JIMT-1 cells inhibited intracellular accumulation of cell surface biotinylated HER2 when compared to control transfected cells (Fig. 3h). Furthermore, a biochemical recycling assay, in which biotin is cleaved off from an internalized biotinylated receptor upon recycling back to the plasma membrane, revealed that SORLA-GFP-expressing JIMT-1 cells had a higher rate of HER2 recycling when compared to JIMT-1 control cells (Fig. 3i). Taken together, these findings demonstrate a new role of SORLA in regulation of HER2 recycling.

### Silencing SORLA triggers HER2 accumulation in dysfunctional lysosomes

As SORLA silencing triggered increased intracellular retention of HER2 in the internalization assays, we wanted to investigate the subcellular localization of HER2 in SORLA-silenced MDA-MB-361 cells. Imaging revealed striking accumulation of HER2 in enlarged LAMP-1-positive structures (late endosomes/lysosomes), not observed in control cells (Fig. 4a, Supplementary Fig. 3a). This unexpected intracellular distribution could be linked to the specific attenuation of PI3K/AKT signaling in SORLA-silenced cells (Fig. 2f, Supplementary Fig. 2g) since the ligand for PI3K, PI(4,5,)P2, is predominantly enriched on the plasma membrane and considered to be absent in late endosomes, thus restricting signaling of the lysosomally localized HER2 through the PI3K/AKT pathway. The accumulation of HER2 in enlarged LAMP-1-positive structures in SORLA-silenced cells was striking as normally lysosomal targeting of growth factor receptors is linked to their rapid degradation. However, on the other hand, HER2 protein levels in SORLA-silenced cells were only modestly reduced (Supplementary Fig. 3a). This apparent discrepancy lead us to investigate whether loss of SORLA could be linked to potentially abnormal lysosome function. Our hypothesis was supported by strong perinuclear accumulation of LAMP1- and CD63 (LAMP3)-positive late endosomes/lysosomes in SORLA-silenced cells compared to control cells (Fig. 4b, c, Supplementary Fig 3b, c). Lysosomal aggregation was confirmed with four different siRNAs targeting SORLA in both MDA-MB-361 and BT474 cells (Supplementary Fig. 3d). Thus, depletion of endogenous SORLA in breast cancer cells leads to altered subcellular localization of lysosomes.

**Figure 4.**
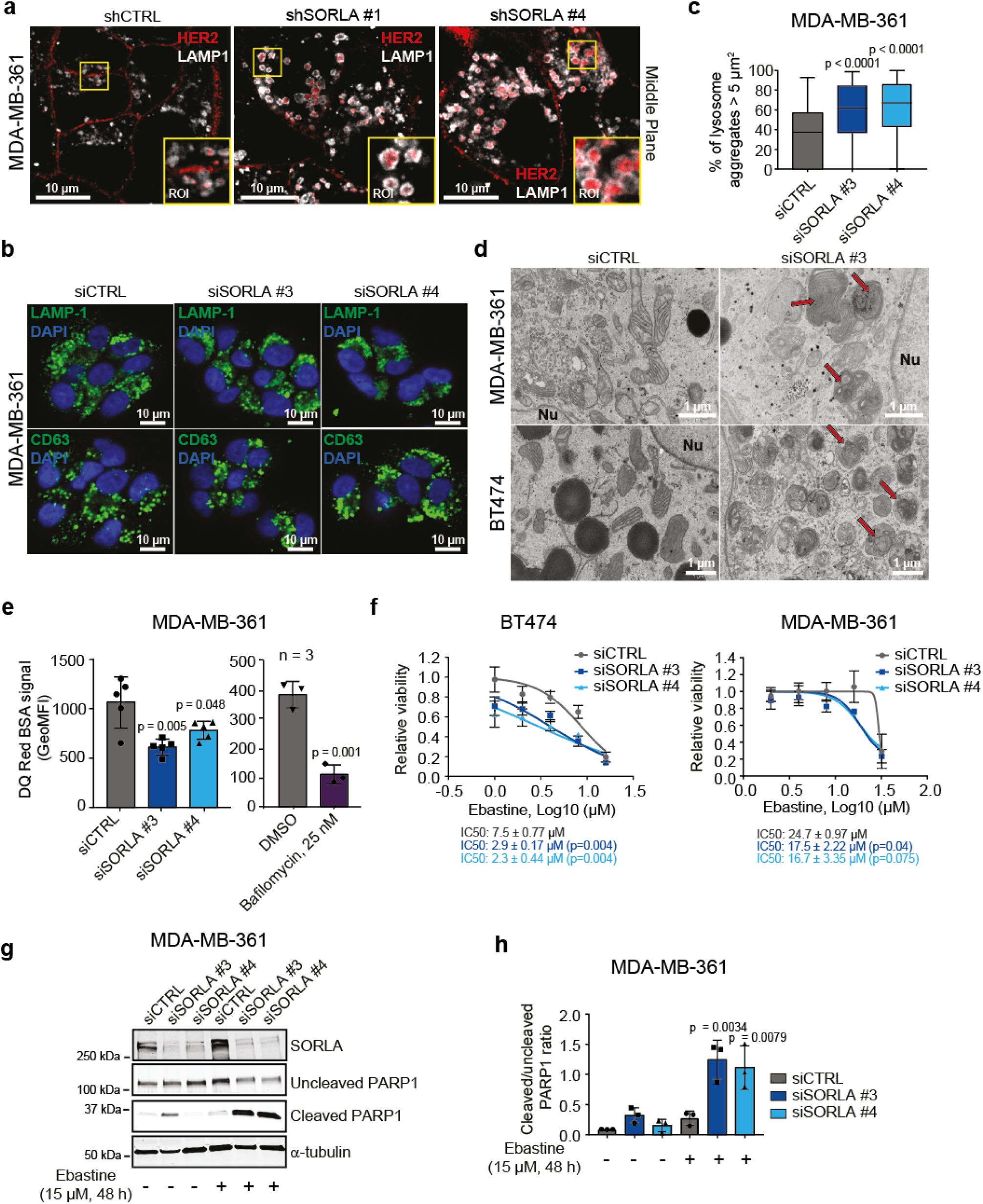
Silencing SORLA induces HER2 accumulation in dysfunctional lysosomes. (**a**) Microscopy images of LAMP1 (white) and HER2 (red) in scramble (shCTRL) and SORLA (shSORLA #1 and shSORLA #4) silenced MDA-MB-361 cells. (**b**) Immunofluorescence imaging of LAMP1 (green) and CD63 (LAMP3; green) in MDA-MB-361 cells (blue is DAPI) after scramble (siCTRL) or SORLA (siSORLA #3 and siSORLA #4) siRNA silencing. (**c**) Quantification of late-endosomes/lysosome aggregation after SORLA silencing in MDA-MB-361 cells. LAMP1-positive structures ≥ 5 µm^2^ were considered as lysosome aggregates (data are displayed as box plots; n = 67-79 cells; statistical analysis: Mann-Whitney test). (**d**) Transmission electron microscopy imaging of lysosomes after scramble (siCTRL) or SORLA (siSORLA #3) silencing in MDA-MB-361 and BT474 cells. Red arrow indicates the maturation defect in late endosome/lysosome structures. (**e**) Flow cytometry analysis of fluorescence signal in 24 h DQ Red BSA-loaded MDA-MB-361 cells after scramble (siCTRL) or SORLA (siSORLA #3 and siSORLA #4) silencing. Cells loaded with DQ Red BSA for 4 h and treated with bafilomycin (or vehicle) treatment are included as controls (bafilomycin blocks lysosome function). Data are mean ± s.d. (n = 5 independent experiments; statistical analysis: unpaired Student’s *t*-test). (**f**) Cell viability assay to determine ebastine (48 h treatment) IC50 values for SORLA- or control-silenced BT474 and MDA-MB-361 cells. Data are mean ± s.d. (n = 12 (4 technical replicates, 3 independent experiments); statistical analysis: unpaired Student’s *t*-test). (**g**) Immunoblotting of cleaved PARP1 from ebastine-treated (15 µM, 48 h), scramble (siCTRL) and SORLA (siSORLA #3 and siSORLA #4) silenced MDA-MB-361 cells. α-tubulin is a loading control. (**h**) Quantification of the ratio of cleaved to uncleaved PARP1 signal from (g). Data are mean ± s.d. (n = 3 independent experiments; statistical analysis: unpaired Student’s t-test). Box plots represent median and 25^th^ and 75^th^ percentiles (interquartile range; IQR) and whiskers extend to maximum and minimum values. Where micrographs are shown, these are representative of n = 3-5 independent experiments. Nu: nucleus.

Next we wanted to investigate if the observed lysosome phenotype is linked to decreased HER2 function. Silencing HER2 with two individual siRNAs affected lysosome localization similarly, but to a lesser extent, than SORLA silencing in BT474 and MDA-MB-361 cells (Supplementary Fig 3e, f). When SORLA was silenced in non-HER2-amplified FGFR2-amplified MFM-223 breast cancer cells (Fig. 1a), there was no effect on lysosomal distribution/localization (Supplementary Fig. 3g). Thus, the lysosome phenotype seems to be linked to the absence of SORLA or HER function in HER2-dependent cells and to impaired cell growth.

Further analyses using transmission electron microscopy (TEM) revealed enlarged lysosomes in SORLA-silenced cells suggesting potential lysosome maturation defects (Fig. 4d). To monitor the proteolytic activity of lysosomes, we analyzed loss of quenching of a fluorogenic protease substrate, DQ Red BSA, loaded into cells ^33^. SORLA-silenced MDA-MB-361 cells showed significantly lower DQ Red BSA signal (indicating reduced lysosomal cleavage of BSA), detected either by confocal microscopy imaging (Supplementary Fig 3h) or by flow cytometry (Fig. 4e), than the respective control cells. These data together indicate a link between SORLA-dependent HER2 signaling and lysosome integrity in HER2-driven cancer cells.

### Depletion of SORLA renders HER2-driven cancer cells sensitive to CADs

Previous studies indicate that cancer cells possess functionally abnormal lysosomes making them more susceptible to cationic amphiphilic drugs (CADs), a heterogeneous class of molecules with a similar chemical structure resulting in lysosomal accumulation and increased lysosomal membrane permeabilization ^34^. Recently, cancer cells were shown to be more sensitive to CAD-induced cell death than non-transformed cells ^35^. Given the defective lysosome integrity of SORLA-depleted cells, we wanted to evaluate the response of anti-HER2 therapy sensitive BT474 and therapy-resistant MDA-MB-361 cells to antihistamine ebastine, a CAD with cytotoxic effects in lung cancer ^35^. Interestingly, the ebastine IC50 values, for inhibiting cell viability, were significantly lower in both of these HER2-amplified breast cancer cell lines when SORLA was silenced compared to control cells (Fig. 4f). Moreover, treatment of SORLA-silenced MDA-MB-361 cells with 15 µM ebastine, which is close to the determined IC50 value that inhibits the growth of these cells, significantly increased levels of cleaved PARP1, indicative of apoptosis, whereas no such effect was seen in control-silenced cells (Fig. 4g,h). Thus, deletion of SORLA increased the sensitivity of anti-HER2 therapy sensitive and resistant breast cancer cells to CADs, presumably due to apoptosis triggered by lysosomal dysfunction. These data indicate that compromised lysosomal integrity of SORLA-silenced cells could be exploited therapeutically to induce cell death of HER2-therapy resistant breast cancer cells.

### SORLA correlates with HER2 and regulates proliferation and tumor growth in bladder cancer

Our *in vitro* data and observations of reduced tumor formation in mice demonstrate an important role for SORLA in regulation of HER2 function in breast cancer cells. To broaden our study for other cancer types in which HER2 overexpression is common and clinically relevant ^36, 37^, we stained SORLA in a bladder cancer tissue microarray (TMA) of 199 patients. In this cancer type HER2 and SORLA levels correlated significantly (Chi-square test. p = 0.0092), whereas there was no correlation between SORLA and EGFR (Fig. 5a, Supplementary Fig. 4a). This led us to test if SORLA also plays a role in the regulation of bladder cancer cell proliferation and *in vivo* tumor growth. Silencing of SORLA in 5637 cells, a bladder carcinoma cell line with a HER2 signaling activating mutation ^31^, significantly inhibited their proliferation (Fig. 5b,c). Moreover, subcutaneous grafting of transiently SORLA-silenced 5637 cells (the strong anti-proliferative effect of shSORLA in vitro precluded sufficient propagation of stably silenced cells for in vivo experiments), in Nude mice resulted in impaired tumor growth (mean tumor volumes of 78.7 mm^3^ and 47.7 mm^3^ (p=0.0461) with control and SORLA-silenced cells, respectively) (Fig. 5d, Supplementary Fig. 4b).

**Figure 5.**
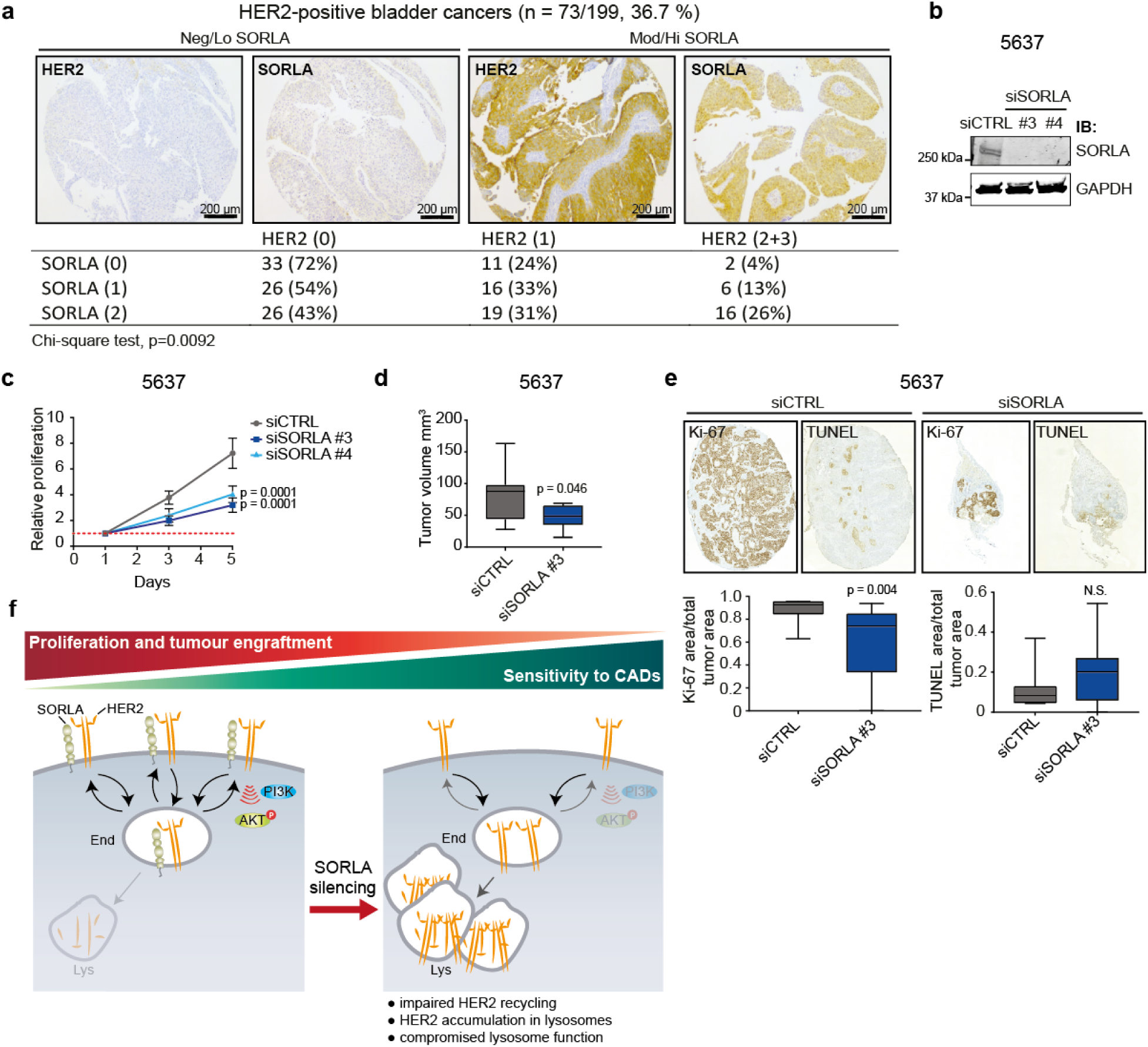
SORLA expression correlates with HER2 and promotes tumorigenesis in bladder carcinoma. (**a**) Immunohistological staining of TMA material and correlation analyses of SORLA and HER2 in clinical bladder cancer specimens. Numbers indicate staining intensity, 0=negative, 1=weak, 2=moderate, 3=high. (**b**) Western blot analysis of SORLA levels in scramble (siCTRL) and siSORLA (siSORLA#3, siSORLA#4) silenced 5637 bladder cancer cells. GAPDH is a loading control. (**c**) Proliferation of scramble (siCTRL) and siSORLA (siSORLA#3, siSORLA#4) silenced 5637 cells. Proliferation data are mean ± s.d. (n = 4 independent experiments; statistical analysis: two-way ANOVA). (**d**) Analysis of tumor growth of subcutaneously injected 5637 cells, with transient SORLA (siSORLA #3) or scramble (siCTRL) silencing, at day 29 in nude mice. Data are displayed as box plots (n = 9-10 mice/group; statistical analysis: unpaired Student’s *t*-test). (**e**) Ki-67 and TUNEL staining of tumor samples prepared as described in (d) and quantifications displayed as box plots (n = 9-10 mice/group; statistical analysis: Mann-Whitney test). (**f**) Schematic illustrating the role of SORLA in regulating the oncogenic fitness of HER2-driven cancer cells. SORLA associates with HER2 via its extracellular domain, co-traffics with HER2 and facilitates HER2 recycling to the plasma membrane to support HER2 downstream signaling. In the absence of SORLA, HER2 becomes localized to enlarged, partially dysfunctional lysosomes resulting in defective HER2 signaling and increased sensitivity to CADs such as ebastine. End: endosome, Lys: lysosome. Box plots represent median and 25^th^ and 75^th^ percentiles (interquartile range; IQR) and whiskers extend to maximum and minimum values.

Furthermore, SORLA-silenced 5637 tumors showed decreased proliferation, but not significantly increased apoptosis (Fig. 5e). These results demonstrate that SORLA-mediated regulation of proliferation is not only restricted to HER2-amplified breast cancers, but is biologically important at least in HER2-driven urothelial cancer. It may also be relevant to other neoplasms considering that data mining of the TCGA database revealed a significant positive correlation between *ErbB2* and *SORL1* expression in testicular germ cell tumors, cervical squamous cell carcinoma and endocervical adenocarcinoma, kidney renal clear cell carcinoma, sarcoma and thymoma (Supplementary Fig. 4c).

## Discussion

Here we demonstrate that SORLA, a sorting protein previously not investigated in carcinomas, is highly expressed in many HER2-driven cancer cell lines, and that SORLA regulates HER2 subcellular localization by associating with the receptor and coupling it to the recycling machinery (Fig. 5f). We find that HER2 distribution between the plasma membrane and endosomes is highly heterogeneous in cancer cells and those with lower SORLA expression exhibit a trend of harboring a substantial pool of intracellular HER2 at steady state. Furthermore, silencing of SORLA induces intracellular accumulation of HER2, and overexpressing SORLA triggers cell-surface localization of HER2. Thus, HER2 undergoes rapid endosomal traffic and recycling, the kinetics of which is regulated by SORLA-HER2 association. The fact that interfering with SORLA 1) dramatically reduced proliferation of HER2-driven bladder and breast cancer cells *in vitro* and *in vivo*, 2) altered HER2 signaling, 3) interfered with lysosome integrity and 4) had additive pro-apoptotic effects in combination with a clinically well-tolerated and widely used lysosome accumulating drug provides new insight into the pathophysiology and targetability of HER2-driven oncogenesis.

Previous studies have shown discrepant results regarding HER2 trafficking. While some studies have shown that HER2 is resistant to internalization ^13, 14, 32^, others have suggested rapid recycling of HER2 back to plasma membrane ^19^. In both scenarios, HER2 is mainly restricted to the plasma membrane where it can associate with signaling platforms to drive the proliferation and tumorigenesis of cancer cells. Here we investigated the localization of HER2 in six different HER2-amplified breast cancer cell lines and found very different patterns of localization. Our observation of HER2 overlapping with EEA-1- and VPS35-positive endosomes in some of the cell lines, suggests that at least a pool of HER2 moves back to the plasma membrane via retromer-dependent vesicle trafficking ^38^. The heterogeneity in the subcellular localization of HER2 is interesting, since it might reflect the functions of HER2 and the efficiency of therapeutic targeting of HER2. Anti-HER2 therapy (lapatinib, trastuzumab) resistant cells, JIMT-1, HCC1954 and MDA-MB-361 cell lines, displayed more intracellular HER2 compared to the therapy sensitive BT474, HCC1419 and SKBR3 cell lines with predominantly plasma membrane localized HER2. It is an intriguing possibility that the reduced viability of some HER2-dependent cancer cell lines upon depletion of trafficking proteins, such as VPS35, Rab7 and LAMP1 ^39^ might be related to alterations in the trafficking of HER2.

The sorting functions of SORLA have been implicated in Alzheimer’s disease and obesity ^24-26, 30^. We found that the ECD of SORLA associates with HER2, while the intracellular domain of SORLA is required for the correct endosomal and plasma membrane localization of SORLA in cancer cells. Importantly, re-expression of neither ECD-TM nor TM-CD truncation mutants of SORLA were sufficient to support proliferation of SORLA-silenced cells. This suggests that both the ECD association with HER2 and the correct subcellular localization of the SORLA-HER2 complex are necessary for proper SORLA function in supporting HER2-driven oncogenesis.

Lysosomal function is strongly linked to cellular fitness and it is especially important for the nutrient balance in rapidly growing cancer cells. One important finding of our work is that SORLA plays a major and unanticipated role in maintenance of lysosome function in HER2-dependent cancer cells, but not in the FGFR2-amplified, SORLA-positive MFM-223 cancer cells. The effects of SORLA depletion on the lysosome function were two-fold. Lysosomes in SORLA-silenced cells showed perinuclear clustering. This was phenocopied to some extent by HER2 silencing in the same cells, in line with previous reports ^40, 41^. In addition, we found that lysosomes were enlarged, displayed an abnormal maturation defect-like appearance and had reduced proteolytic activity in SORLA-depleted cells. Strikingly, after SORLA silencing, HER2 accumulated into lysosomes without being efficiently degraded (Fig. 5f). This is in stark contrast to the rapid lysosomal degradation of HER2 following treatment with geldanamycin, which inhibits HSP90-CDC37 complex mediated stabilization of HER2 on the plasma membrane ^42, 43^ and suggests that SORLA regulates HER2 in a fundamentally distinct way than HSP90. Possibly due to the mistargeted localization and lack of suitable signaling co-factors on late endosomes, HER2 signaling to the PI3K/AKT/mTOR proliferative pathway was decreased in SORLA-silenced cells. However, lysosomal aggregation triggered by HER2 silencing was not as dramatic as that induced by SORLA silencing. To us this implies that SORLA also contributes to lysosome function via HER2-independent mechanisms. The possible role of SORLA-dependent alteration of lipid and glucose metabolism ^27–30^ in lysosomal dysfunction remains to be investigated.

We found that SORLA-silenced cells undergo apoptosis when exposed to low doses of the antihistamine ebastine, which belongs to a heterogeneous group of molecules, CADs, with similar chemical features ^44^ and with a tendency to accumulate strongly to leaky lysosomes that are common in transformed cells ^34, 35^. CADs, including ebastine, have been utilized in several *in vitro*, pre-clinical and clinical trials to target the vulnerability of lysosomes in cancer ^35^. Importantly, we found that both anti-HER2 therapy resistant and sensitive HER2-amplified breast cancer cells are sensitive to the combination of SORLA silencing and low doses of ebastine. Silencing SORLA alone mainly reduced proliferation, but when combined with ebastine triggered enhanced apoptosis. This additive effect could potentially be exploited by combining current anti-HER2 therapies with CADs, since breast cancer cells and patient-derived xenografts resistant to anti-HER2 drug-antibody conjugate were recently shown to possess dysfunctional lysosomes ^45^.

In conclusion, we have discovered that SORLA regulates the subcellular localization of HER2 and that dynamic recycling is essential for the oncogenic fitness of HER2. The endosomal trafficking of HER2 could provide new rationale for designing novel targeted therapies and understanding the resistance mechanisms induced by the current HER2-targeting therapies.

## Methods

Methods are included in the supplementary information and any relevant references are provided in the reference list of the manuscript

## Author Contributions

Conceptualization, M.P., J.I. and P.S.; Methodology, M.P., J.I., P.S., E.P., P.T., P.B., and O.A.; Investigation, M.P., J.I., P.S., E.P., I.P., N.J., A.P., I.S., and M.B.; Writing – original draft, M.P. and J.I.; Writing – Review&Editing, M.P., J.I. and P.S. Funding acquisition, M.P. and J.I.; Supervision, M.P., J.I., and P.T.

## Acknowledgements

We thank J. Jukkala, P. Laasola and J. Siivonen for technical assistance, M. Saari for help with the microscopes. The Cell Imaging Core, supported by the University of Turku and Biocenter Finland at Turku Centre for Biotechnology for technical assistance with imaging and Turku Centre for Disease modeling, University of Turku for assistance with the animal experiments. H. Hamidi for the scientific illustrations and manuscript editing, J. Westermarck, M. Salmi and the Ivaska lab for critical reading and feed-back on the manuscript. This study has been supported by the Academy of Finland (M.P. and J.I.), Academy of Finland CoE for Translational Cancer Research (J.I. H.J.), ERC CoG grant 615258, Sigrid Juselius Foundation, Orion Research Foundation and the Finnish Cancer Organization (J.I.). P.S. has been supported by the Turku Doctoral Program of Molecular medicine (TuDMM).

## Supplementary video titles

**Supplementary Video 1** – Live-cell TIRF plane imaging of SORLA-GFP and AlexaFluor-568-labeled trastuzumab (Tz-568; red) in MDA-MB-361 cells.

**Supplementary Video 2** – Live-cell confocal imaging of SORLA-GFP and AlexaFluor-568-labeled trastuzumab (Tz-568; red) in the cytoplasm of MDA-MB-361 cells.

---

**Supplementary Figure 1.**
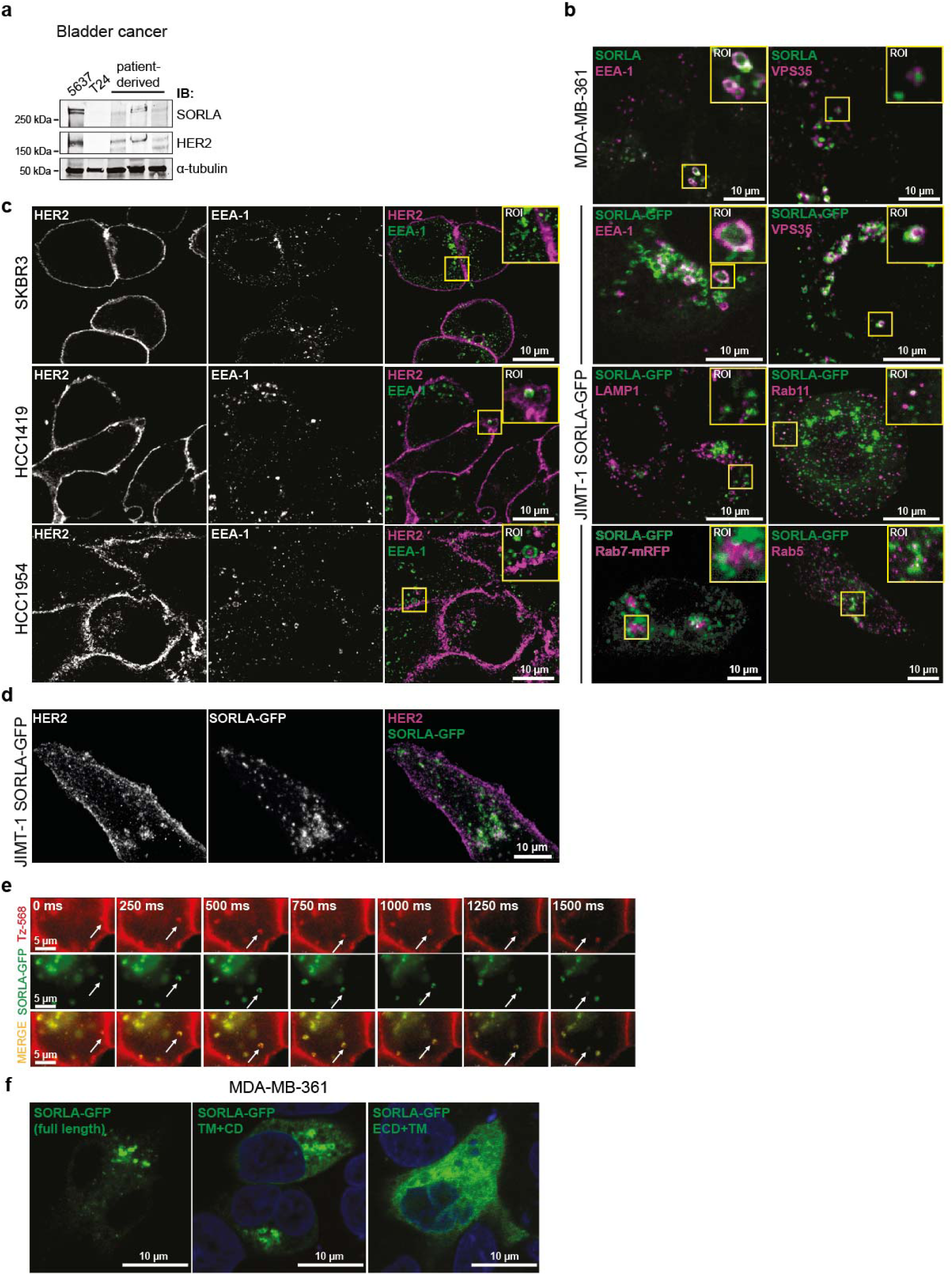
(**a**) Western blot analysis of SORLA and HER2 in bladder cancer cell lines (5637 and T24) and three patient-derived cell cultures. α-tubulin is a loading control. (**b**) Confocal microscopy imaging of endogenous SORLA or SORLA-GFP (green) with endogenous EEA1, VPS35, LAMP1, Rab11, Rab5 (magenta) or Rab7-mRFP (magenta; Rab7 was overexpressed due to lack of reliable Rab7 antibodies for staining) in MDA-MB-361 (top panel) or JIMT-1 cells. (**c**) Endogenous HER2 (magenta) and EEA1 (green) staining in SKBR3, HCC1419 and HCC1954 cells. (**d**) Co-localization between SORLA-GFP and HER2 in JIMT-1 cells (n = 130 cells from 7 independent experiments). Details of colocalization analysis can be found in the methods section. (**e**) Live-cell confocal imaging of AlexaFluor-568-labeled trastuzumab (Tz-568; red) and SORLA-GFP (green) in the cytoplasm of MDA-MB-361 cells, 250 ms frames. (**f**) Confocal imaging of different SORLA-GFP fusion proteins in MDA-MB-361 cells. ECD: extracellular domain; TM: transmembrane domain; CD: cytosolic domain.

**Supplementary Figure 2.**
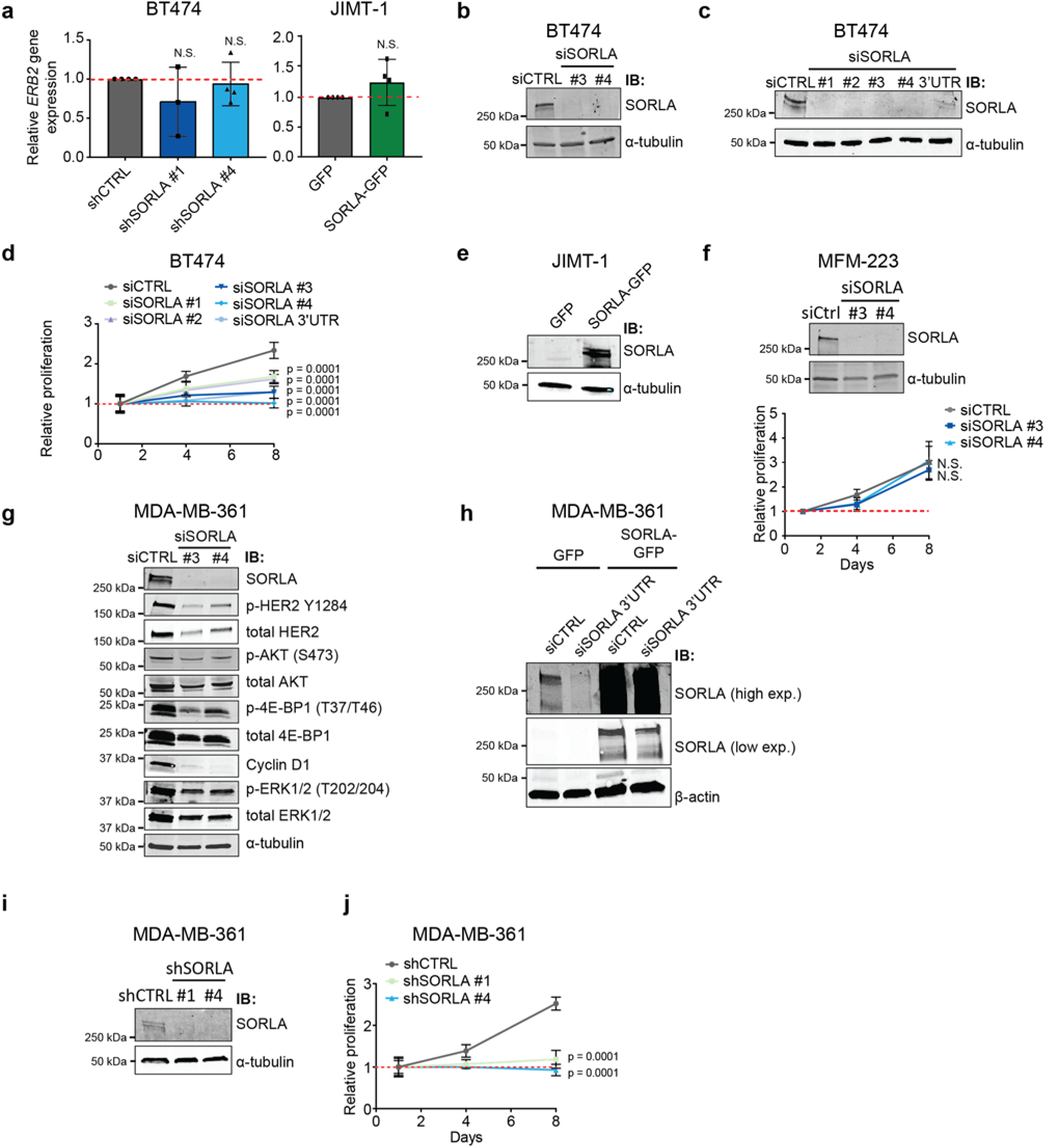
(**a**) Quantitative real-time PCR analysis of *ErbB2* mRNA gene expression levels in the indicated transfectants. Data are mean ± s.d. (n = 4 independent experiments; statistical analysis: unpaired student’s t-test). (**b-c**) Western blot analysis of SORLA in scramble (siCTRL) and SORLA (siSORLA #1, siSORLA #2, siSORLA #3, siSORLA #4, siSORLA 3´UTR against the 3’UTR of SORLA) silenced BT474 cells. α-tubulin is a loading control. (**d**) Proliferation of BT474 cells after SORLA silencing as in (c). Data are mean ± s.d (n = 4 independent experiments; statistical analysis: two-way ANOVA). (**e**) Western blot analysis of SORLA in JIMT-1-GFP and JIMT-1-SORLA-GFP cells. α-tubulin is a loading control. (**f**) Proliferation of scramble (siCTRL) and siSORLA (siSORLA#3, siSORLA#4) silenced non-HER2-amplified MFM-223 breast cancer cells. Proliferation data are mean ± s.d. (n = 12 (4 replicates, 3 independent experiments); statistical analysis: two-way ANOVA). (**g**) Western blot analysis of the indicated signaling proteins (phosphorylated and total) in MDA-MB-361 cells after scramble (siCTRL) and SORLA silencing (siSORLA #3 and siSORLA #4), α-tubulin is a loading control. Shown are representative immunoblots from n = 3 independent experiments. (**h**) Western blot analysis of SORLA in scramble (siCTRL) and SORLA (siSORLA 3´UTR against the 3’UTR) silenced MDA-MB-361-GFP and MDA-MB-361-SORLA-GFP cells. Shown are two exposures for SORLA. β-actin is a loading control. (**i**) Western blot analysis of SORLA in scramble (siCTRL) and shSORLA (shSORLA #1, shSORLA #4) silenced MDA-MB-361 cells. α-tubulin is a loading control. (**j**) Proliferation of MDA-MB-361 shRNA cells silenced as in (i). Data are mean ± s.d. (n = 4 independent experiments; statistical analysis: two-way ANOVA). IB: immunoblotting; high exp: high exposure; low exp: low exposure.

**Supplementary Figure 3.**
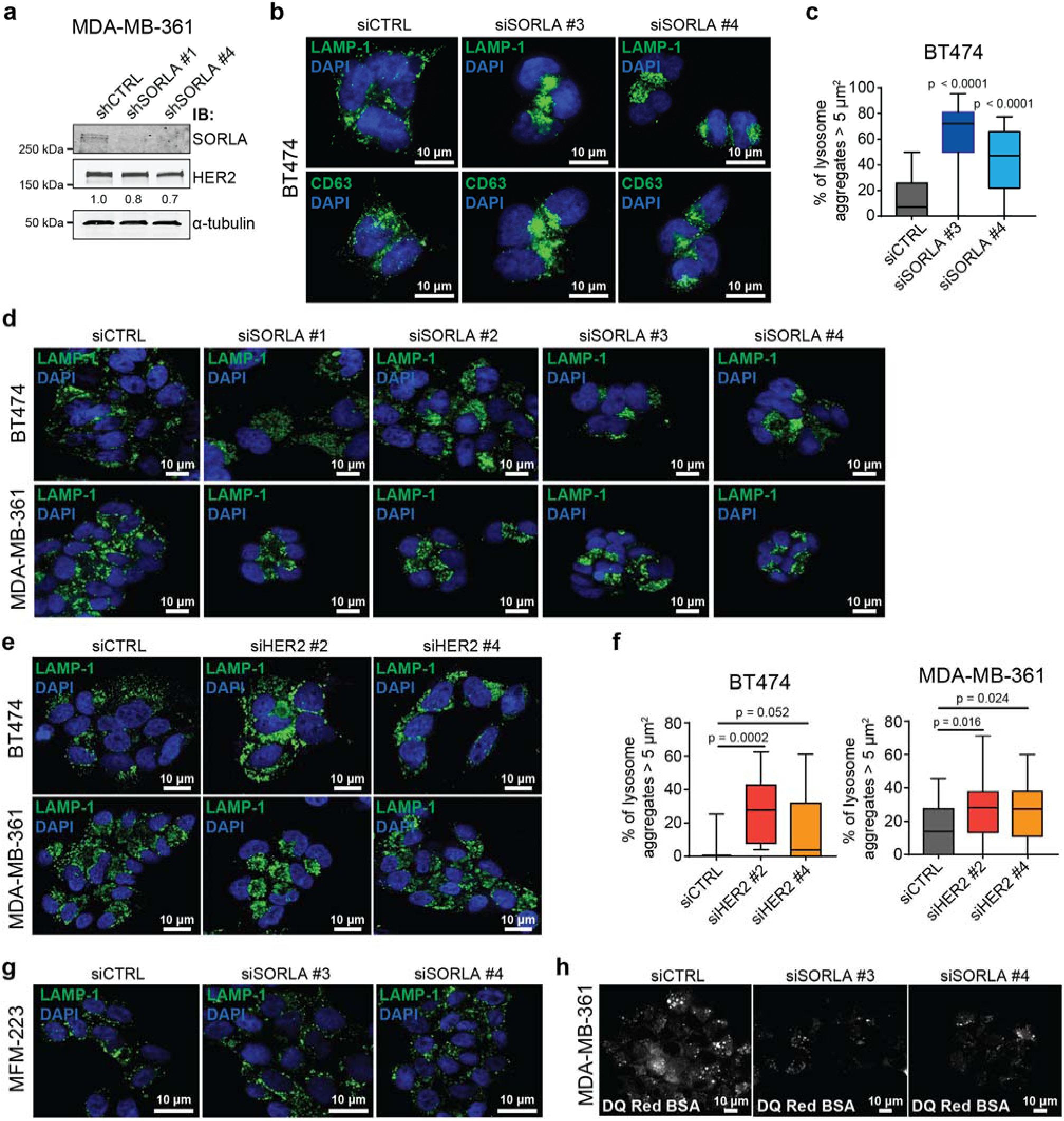
(**a**) Western blot analysis of SORLA and HER2 in scramble (siCTRL) and shSORLA (shSORLA #1, shSORLA #4) silenced MDA-MB-361 cells. α-tubulin is a loading control. Numbers are HER2 levels relative to CTRL. (**b**) Confocal microscopy images of LAMP1, CD63 and DAPI (for nucleus) in scramble (siCTRL) and siSORLA (siSORLA#3, siSORLA#4) silenced BT474 cells. (**c**) Quantification of late-endosomes/lysosome aggregation after SORLA silencing in BT474 cells. LAMP1-positive structures ≥ 5 µm^2^ were considered as lysosome aggregate. Data are displayed as box plots (n = 24-35 cells, three independent experiments; statistical analysis: Mann-Whitney test). (**d**) Confocal microscopy images of LAMP1 and DAPI (for nucleus) in scramble (siCTRL) and siSORLA (siSORLA #1, siSORLA #2, siSORLA#3, siSORLA#4) silenced BT474 and MDA-MB-361 cells. (**e-f**) Immunofluorescence staining of LAMP1 and quantification of late-endosomes/lysosome aggregation after HER2 silencing (and control siRNA) in BT474 and MDA-MB-361 cells. LAMP1-positive structures ≥ 5 µm^2^ were considered as lysosome aggregates (data are displayed as box plots; n = 7-24 BT474 cells and n = 10-34 MDA-MB-361 cells from 2 independent experiments; statistical analysis: Mann-Whitney test). (**g**) Confocal microscopy images of LAMP1 and DAPI (for nucleus) in scramble (siCTRL) and siSORLA (siSORLA#3, siSORLA#4) silenced non-HER2-amplified MFM-223 breast cancer cells. (**f**) Microscopy images of DQ Red BSA degradation (loss of quenching) in 48 h BSA-loaded MDA-MB-361 cells after scramble (siCTRL) or SORLA (siSORLA #3 and siSORLA #4) silencing. Box plots represent median and 25^th^ and 75^th^ percentiles (interquartile range; IQR) and whiskers extend to maximum and minimum values. IB: immunoblotting.

**Supplementary Figure 4.**
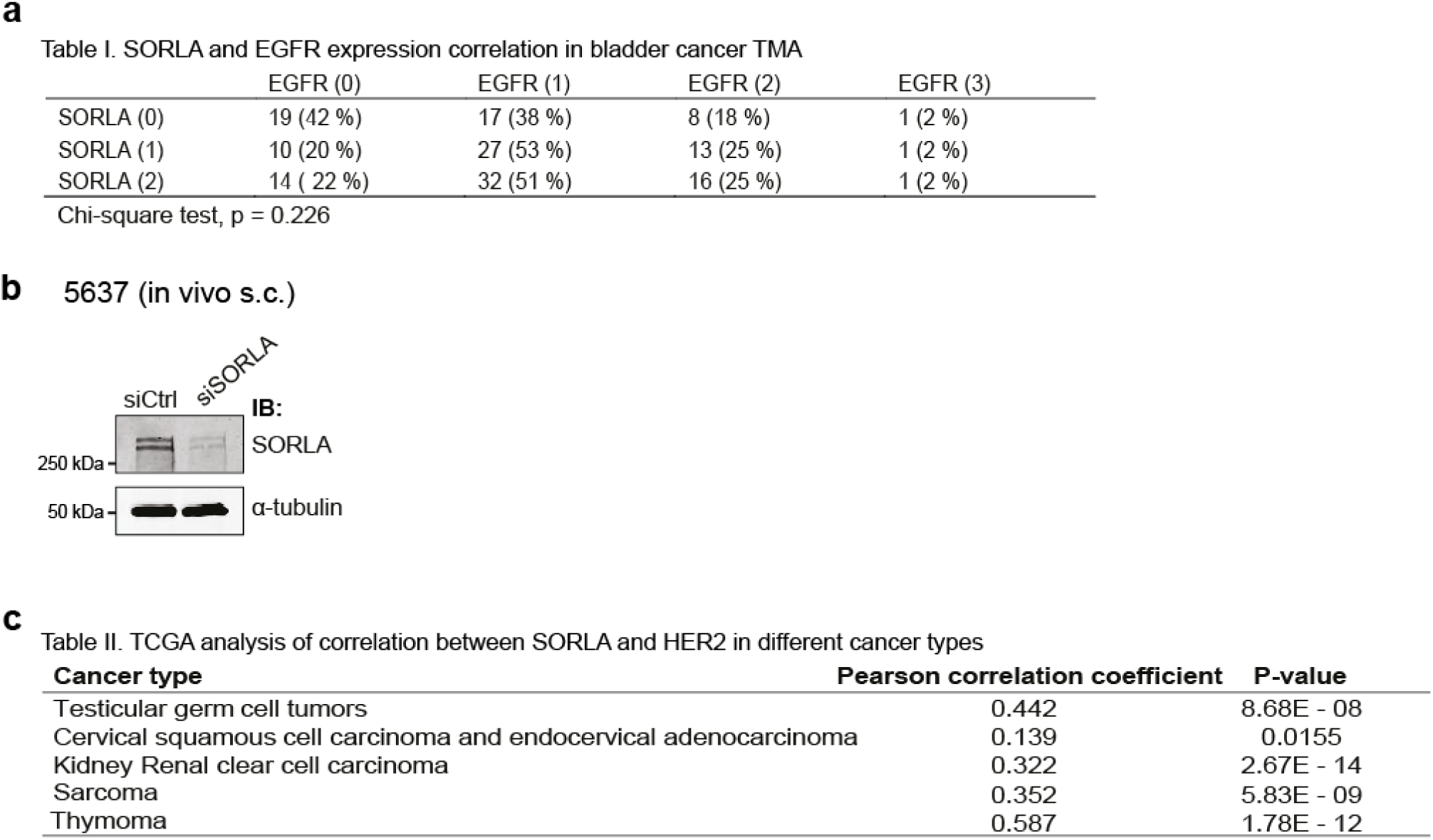
(**a**) Correlation analyses of SORLA and EGFR expression in bladder cancer TMA material. Numbers indicate staining intensity, 0=negative. (**b**) Western blot analysis of SORLA levels in scramble (siCTRL) and siSORLA (siSORLA#3) silenced 5637 bladder cancer cells used in the in vivo xenograft assay in Fig. 5d,e. α-tubulin is a loading control. (**c**) Correlation between SORLA and HER2 expression in different neoplasms based on TCGA data.

## Methods

### Subcutaneous and Ductal carcinoma in situ (DCIS) in vivo xenograft models

For subcutaneous (s.c.) tumors, 2 million siCTRL and siSORLA #3 treated 5637 bladder cancer cells were injected s.c. in 100 µl (50% Matrigel, 50% PBS) at the flank of 6 weeks old Nude mice. Mice were sacrificed after 29 days, and tumors were dissected, fixed in 10% formalin, and processed for paraffin sections with standard protocols. Sections were stained with hematoxylin-eosin (HE) and immunohistochemistry (IHC) for proliferation (Ki67) and apoptosis (TUNEL) markers. Sections were imaged with pannoramic slide scanner.

For intraductal tumor xenografts, MDA-MB-361 shCTRL and shSORLA-silenced breast cancer cells were resuspended in PBS (25 000 cells/ul). Trypan blue (0.1%) was added to the cell solution to visualize successful injection. Eight to ten weeks old female NOD.SCID mice were medicated with Temgesic for analgesia and anesthetized with isoflurane. After removal of abdominal hair, the tit of the abdominal (4^th^) mammary glands was carefully snipped, and 4 µl (100 000 cells) of cell suspension was injected into the mammary ducts. A 30G Hamilton syringe with 50-μl capacity and a blunt-ended needle was used for the injection under a stereomicroscope. Post-operatively, the mice were further dosed with Rimadyl for pain relief. Mice were sacrificed 10 weeks after tumor inoculation, and abdominal mammary glands were dissected. Mammary gland whole mounts were prepared on object glasses and fixed in Carnoy´s medium (60% EtOH, 30% chloroform, 10% glacial acetic acid) overnight (o/n) at +4°C. After rehydration in decreasing EtOH series and staining with carmine alum (0.2% carmine, 0.5% aluminium potassium sulphate dodecahydrate) o/n at room temperature (RT), samples were dehydrated and cleared in xylene for 2-3 days. Samples were mounted in DPX Mountant for histology (Sigma) and images were taken with Zeiss SteREO Lumar V12 stereomicroscope (NeoLumar 0.8× objective, Zeiss AxioCam ICc3 colour camera). All images per gland were combined automatically into a mosaic picture with PhotoShop. DCIS area per gland was quantified in ImageJ.

All animal studies were ethically performed and authorized by the National Animal Experiment Board and in accordance with The Finnish Act on Animal Experimentation (Animal license number ESAVI-9339-04.10.07-2016).

### Cell lines and cell culture

MDA-MB-361 cells (ATCC) were grown in Dulbecco’s modified essential medium (DMEM; Sigma-Aldrich, D5769) supplemented with 20 % fetal bovine serum (FBS; Sigma-Aldrich, F7524), 1 % vol/vol Penicillin/Streptomycin (Sigma-Aldrich, P0781-100ML) and L-glutamine. BT474 and 5637 cells (from ATCC) were grown in RPMI1640 (Sigma-Aldrich, R5886) supplemented with 10 % FBS, 1 % vol/vol Penicillin/Streptomycin and L-glutamine. JIMT-1 (obtained from DMZK), HCC1954 (ATCC), HCC1419 (ATCC), MCF7 (ATCC), MDA-MB-231 (ATCC), MDA-MB-436 (ATCC) and MFM-223 (ATCC) were grown in DMEM supplemented with 10 % FBS, 1 % Penicillin/Streptomycin and L-glutamine. MCF10A (ATCC) and MCF10A DCIS.com (provided by J.F. Marshall (Barts Cancer Institute, Queen Mary University of London, London, England, UK)) were grown in DMEM/F12 (Invitrogen, #11330-032) supplemented with 5 % horse serum (Invitrogen#16050-122), 20 ng/ml human epidermal growth factor (Sigma-Aldrich, E9644), 0.5 mg/ml hydrocortisone (H0888-1G; Sigma-Aldrich), 100 ng/ml Insulin (Sigma-Aldrich, I9278-5ML) and 1 % vol/vol Pen/Strep. All cells were regularly tested for mycoplasma infection and were grown at +37°C, 5% CO_2_, until 70-80 % confluency before detaching and plating into new dish. Media was changed every three days.

### Antibodies

The antibodies used are described in supplementary table 1.

### Generation of lentiviral shRNA and SORLA-GFP particles

Lentiviral particles containing sequences encoding shRNA SORLA and GFP or control scramble sequence and GFP or particles encoding for SORLA-GFP or GFP alone were generated in 293FT packaging cell line (complete medium: high glucose DMEM, 10% FBS, 0.1mM NEAA, 1mM MEM Sodium Pyruvate, 6mM L-Glutamine, 1% Pen/Strep and 0.5mg/ml Geneticin) by transient transfection of transfer vector (#TL309181A SORL1 SR031650, #TL309181B SORL1 SR031650, # TL309181C SORL1 SR031650, TL309181D SORL1 SR031650, or scramble control (#TR30021, Origene), 2^nd^ generation packaging plasmid-psPAX2 (Addgene #12259) and envelope vector-pMD2 (Addgene #12260) with the ratio (7:2:1) using calcium-phosphate precipitation method (Graham FL, 1973). 72h post transfection medium containing viral vectors was collected, concentrated for 2h by ultracentrifugation (26,000g) in swing-out rotor SW-32Ti (Beckman Coulter, Brea, US-CA), resuspended in residual medium and flash frozen in liquid nitrogen. Functional titer was evaluated in 293FT cells by FACS (BD LSRFortessa, Becton Dickinson).

### Lentiviral transduction to generate stable cell lines

To generate stable silenced cell lines 4×10^5^ BT-474 or MDA-MB-361 cells were seeded on 10 cm dishes and transduced 24 h later with MOI 47 of lentivirus in low volume of full media. To obtain stable overexpression of SORLA-GFP or GFP 8×10^4^ JIMT-1 or MDA-MB-361 cells were seeded in 24-well plate and transduced 24h later with MOI 60. Medium containing viral particles was removed 16h later. Cells expressing GFP, indicative of lentiviral integration were collected by fluorescence assisted cell sorting (BD FACSaria II cell sorter, Becton Dickinson, Franklin Lakes, US-NJ) with a gating strategy to obtain medium expression.

### Transient transfections

For transient protein expression, Lipofectamine 3000 (Invitrogen, P/N 100022052) and P3000 enhancer reagent (Invitrogen, P/N100022058) were used according to manufacturer’s instructions. Cells were transfected with plasmids 24 hours prior to experiments. Transient siRNA transfections were performed using Lipofectamine RNAiMAX reagent (Invitrogen, P/N 56532) according to manufacturer’s instructions. SORLA targeting siRNAs were ON-TARGETplus obtained from Dharmacon: siSORLA #1 (J-004722-08), siSORLA #2 (J-004722-06), siSORLA #3 (J-004722-07), siSORLA #4 (J-004722-05). For rescue experiments, siRNA against 3’UTR end of SORLA was obtained from Qiagen: siSORLA 3’UTR (SI05039888). HER2 targeting siRNAs were ON-TARGETplus obtained from Dharmacon: siHER2 #2 (J-003126-17), siHER2 #4 (J-003126-20). For controls, Allstars negative control (Qiagen, Cat. No. 1027281) was used. SiRNA concentrations used ranged between 20-40 nM and cells were transfected with siRNAs 48 hours prior to experiments.

### Western blot analysis

Protein extracts were separated using SDS-PAGE under denaturing conditions (4–20% Mini-PROTEAN TGX Gels) and were transferred to the nitrocellulose membrane (Bio-Rad Laboratories). Membranes were blocked with 5% milk-TBST (Tris-buffered saline and 0.1% Tween 20), incubated with the indicated primary antibodies overnight at +4°C. Primary antibodies were diluted in blocking buffer (Thermo, StartingBlock (PBS) blocking, Product#37538) and PBS (1:1 ratio) mix and incubated overnight at +4°C. Primary antibody dilutions used ranged from 1:500 to 1:1000. After primary antibody incubation, membranes were washed three times with TBST and incubated with fluorophore-conjugated secondary antibodies were diluted (1:1000) in the blocking buffer at RT for 1 h. Membranes were scanned using an infrared imaging system (Odyssey; LI-COR Biosciences). The following secondary antibodies were used: donkey anti-mouse IRDye 800CW (LI-COR, 926-32212), donkey anti-mouse IRDye 680RD (LI-COR, 926-68072), donkey anti-rabbit IRDye 800CW (LI-COR, 926-32213), donkey anti-rabbit IRDye 680RD (LI-COR, 926-68073).

### Co-immunoprecipitations

Non-transfected cells or cell transfected as indicated were lysed in IP-lysis buffer (0.5% Triton X-100, 10 mM Pipes, pH 6.8, 150 mM NaCl, 150 mM sucrose, 3 mM MgCl2, complete, and phosphatase inhibitors [Mediq; Roche]), cleared by centrifugation, and incubated with GFP-trap beads to pull-down GFP-proteins (Chromotek; gtak-20) for 1 h at +4°C or with mouse anti-HER2 antibody (1 µg/sample; ThermoScientific, MA5-14057) or isotype matching IgG control antibody at +4 °C overnight. Antibody complexes were bound to 0.5% BSA pre-blocked protein-G sepharose beads for 1 h at 4°C. Complexes bound to the beads were isolated using 1000 g 3 min centrifugation, washed 3 times with washing buffer (20 mM Tris-HCl (pH 7.5), 150 mM NaCl, 1 % NP-40; 500 µl) and eluted in sample buffer. Input and precipitate samples were analyzed by Western blotting. Primary antibodies were incubated overnight at +4°C. Mouse anti-HER2 (ThermoScientific; MA5-14057), rabbit anti-GFP (Molecular Probes; A11122) and mouse anti-SORL1 (BD Transduction Lab; 612633) diluted 1:1000 in 5 % milk in TBST were used followed by the appropriate IRDye conjugated secondary antibodies.

### Immunofluorescence staining and imaging

Cells were plated on µ-Slide 8-well (Ibidi, 80826) or in some cases in µ-dish 3.5 mm dishes (Ibidi, 80136). Cells were fixed with 4 % paraformaldehyde (PFA) 10 min at RT, quenched with 50 mM NH_4_Cl for 15 min at RT, blocked and permeabilized with 30 % horse serum in PBS + 0.3 % Triton X-100 10 min at RT and incubated with primary antibodies diluted in 30 % horse serum overnight at +4°C. Stainings were performed using antibodies against HER2 (Herceptin (Roche) 0.15 μg/ml or mouse monoclonal antibody (ThermoScientific; MA5-14057; dilution of 1:300), LAMP1 (Santa Cruz; SC-20011 (H4A3), dilution 1:50), SORLA (rabbit monoclonal, CM Petersen Lab, Århus University, dilution of 1:300), EEA-1 (goat polyclonal, Santa Cruz; sc-6415, dilution 1:50), VPS35 (goat polyclonal, Abcam; ab10099, dilution 1:300), CD63 (mouse mAb, Hybridoma Bank; H5C6, dilution 1:300) and Rab11 (rabbit polyclonal, Cell Signaling Technology, #5589, dilution 1:100). After washings appropriate secondary antibodies (donkey anti-mouse AlexaFluor 488 (LifeTechnologies, A21202), donkey anti-rabbit AlexaFluor 488 (Invitrogen,), goat anti-human AlexaFluor 568 (Invitrogen, A21090), donkey anti-goat AlexaFluor 647 (Invitrogen, A21447), donkey anti-mouse AlexaFluor (Invitrogen, A31571)) diluted 1:300 in 30 % horse serum were added together with DAPI (1:1000) for 1 hour at room temperature. After PBS washings the samples were imaged right away or stored at +4°C in dark. Imaging was performed either with Carl Zeiss LSM780 laser scanning confocal microscope or a 3i CSU-W1 spinning disk confocal microscope with Hamamatsu CMOS (63x objective).

### Transmission electron microscopy

The cells were fixed in 5% glutaraldehyde in 0.16 M s-collidine buffer, pH 7.4. The samples were post-fixed for 2 h with 1% OsO4 containing 1.5% potassium ferrocyanide, dehydrated with a series of increasing ethanol concentrations and embedded in 45359 Fluka Epoxy Embedding Medium kit. 70-nm sections were cut with an ultramicrotome, and stained with 1% uranyl acetate and 0.3% lead citrate. The sections were examined with a JEOL JEM-1400 Plus transmission electron microscope.

### Live-cell imaging

Lentiviral transduced SORLA-GFP expressing MDA-MB-361 cells were kept on ice and washed twice with ice cold PBS. Alexa-567-conjugated trastuzumab (0.15 μg/ml) in Hank’s Balanced Salt Solution was incubated with the cells on ice for 1 hr protected from light. The cells were washed twice with ice cold PBS before adding pre-warmed culture media supplemented with 5% HEPES (without serum). Imaging was performed with Deltavision OMX V4 total internal reflection microscopy (TIRFM; GE Healthcare) every 250 ms with 60x/1.49 objective (Olympus TIRF objective).

### Proliferation assay

Cells, transfected as indicated, were plated on 96-well plate (3000 cells/well) in volume of 100 µl. After 1d, 4d and 8d of cell growth, 10 µl/well of WST-8 reagent (cell counting kit 8, Sigma-Aldrich, 96992) was added and absorbance at 450nm was measured by a plate reader (Thermo, Multiscan Ascent) after 1-2 h of incubation at +37°C with 5 % CO_2_. Medium without cells was used as background and the A450 of background was subtracted from the samples. Relative proliferation was calculated by normalizing the A450 values of 4d and 8d to 1d A450 values. For drug sensitivity assays cells were treated the following day after plating with increasing concentrations of ebastine (BT474: 2, 4, 8, 16, 32 µM and MDA-MB-361: 2, 4, 8, 16, 32, 64 µM). DMSO was used as a control and proliferation was measured after 48 h.

### Analysis of cell surface HER2 levels by flow cytometry

Cells were detached by HyQtase, fixed with 4 % PFA (in PBS for 15 min at RT), washed before staining with 1:200 dilution of rabbit anti-HER2 (9G6, Abcam; Ab16899) primary antibody for 1 h at +4°C with rotation. After washing with PBS donkey anti-rabbit AlexaFluor 647 (Invitrogen, A31573) secondary antibody (dilution 1:300 in PBS) was added for 1h at 4°C, cells were washed with PBS and analyzed with LSRFortessa (BD Biosciences). Data analysis was performed with Flowing software version 2 (Cell Imaging Core of the Turku Centre for Biotechnology). The fluorescence intensity geometric mean of cells stained with secondary antibody alone was used as background and subtracted from the stained samples.

### DQ Red BSA assay

For microscopy assays, silenced cells were plated on ibidi 8-well µ-slide 72 hours post-transfection. On the next day, the medium was replaced with DQ Red BSA (ThermoScientific, D12051)-containing media (25 µg/ml, 200 ul/well) and cells were incubated for 48 h at 37°C with 5 % CO_2_, the cells were washed, fixed with 4 % PFA for 10 minutes at room temperature, washed again and imaged with Carl Zeiss LSM780 laser scanning confocal microscope.

For flow cytometric assays, the cells were silenced and loaded with DQ Red BSA (25 µg/ml, 1 ml per well in 6-well plate for 24 h) as above. When indicated, 25 nm bafilomycin (Calbiochem, 196000-10UG) or DMSO only was added to DQ Red BSA and the loading was shortened to 4 h. The loaded cells were detached with HyQtase, washed, fixed with 4 % PFA for 15 min at room temperature, washed, and analyzed using LSRFortessa (BD Biosciences) and Flowing software. Fluorescence signals from unstained cells (background) were subtracted from the DQ Red BSA signals and geometric means were plotted.

### RNA extraction, cDNA synthesis and qPCR

Cells were lysed in RA lysis buffer and RNA was extracted according to manufacturer’s instructions (NucleoSpin RNA extraction kit, Macherey-Nagel, 740955.5). RNA concentrations were measured by NanoDrop Lite (Thermo). Complementary DNA (cDNA) was synthesized using a high-capacity cDNA Reverse Transcription Kit (Applied Biosystems) according to the manufacturer’s instructions. Quantitative real time PCR reactions with TaqMan probes were performed according to manufacturer’s instructions (Thermo/Applied Biosystems, TaqMan™ Universal Master Mix II, 4440040). The following TaqMan probes (ThermoScientific, 4331182) were used: ErbB2 (Hs01001580_m1). Relative quantification of gene expression values were calculated using the ddCt method ^46^

### Biotin-based HER2 endocytosis assay

HER2 endocytosis was measured using a cell-surface biotinylation-based assay as previously described ^47^. Shortly, siSORLA silenced and control silenced MDA-MB-361 cells, as well as GFP-ctrl and SORLA-GFP expressing JIMT-1 cells were grown to 80% confluence, placed on ice, and washed once with cold PBS. Cell-surface proteins were labelled with 0.5 mg/ml of EZ-link cleavable sulfo-NHS-SS-biotin (#21331; Thermo Scientific) in Hanks’ balanced salt solution (H9269; Sigma) for 30 min at 4°C. Unbound biotin was removed by washing with cold Hanks’ balanced salt solution. Thereafter, the cells were allowed to internalize receptors in prewarmed 10% serum-containing medium at +37°C for the indicated times. Internalization was stopped by transferring the cells on ice and adding cold media. The remaining biotin on the cell surface was removed with 60 mM MesNa (63705; sodium 2-mercaptoethanesulfonate: Fluka) in MesNa buffer (50 mM Tris-HCl [pH 8.6], 100 mM NaCl) for 30 min at 4°C, followed by quenching with 100 mM iodoacetamide (IAA, Sigma) for 15 min on ice. To detect the total surface biotinylation, plates were left on ice after the biotin labelling and MesNa treatment was omitted. Cells were then washed with PBS, scraped in lysis buffer (50 mM Tris pH 7.5, 1.5 % Triton X-100, 100 mM NaCl, complete, and phosphatase inhibitors [Mediq; Roche]) at 4°C for 20 min. After clarification by centrifugation (14,000 × g, 10 min, 4°C), HER2 was immunoprecipitated from the supernatants with appropriate antibodies and protein G sepharose beads (17-0618-01; GE Healthcare). The immunoprecipitates were eluted in non-reducing Laemmli sample buffer and subjected to Western blotting as described above. Biotinylated (internalized) HER2 and total receptor levels were detected by immunoblotting with horseradish peroxidase (HRP)-conjugated anti-biotin antibody (#7075; Cell Signaling Technology) and receptor-specific antibodies, respectively. Enhanced chemiluminescence-detected biotin and receptor signals were quantified as integrated densities of protein bands with ImageJ (v. 1.43u), and each biotin signal was normalized to the corresponding receptor and total biotin signals.

### HER2 recycling assay in JIMT-1 SORLA-GFP cells

HER2 recycling was measured using a cell-surface biotinylation-based assay as previously described ^47^. Cell surface proteins were biotinylated and allowed to be internalized as described above for endocytosis assay for 30 min. After the first MesNa/IAA treatment, receptor recycling was triggered by treating the cells with prewarmed media containing 10% FBS at +37°C for the indicated time followed with a second MesNa/IAA treatment cleaving biotin from cell surface recycled proteins. HER2 was immunoprecipitated and analysed for biotinylation and total levels as in the endocytosis assay.

### Imaging-based trastuzumab internalization assay

Silenced MDA-MB-361 cells were plated on ibidi 35 mm µ-dishes after 72 h of silencing. The following day, cells were washed once with PBS and incubated with Alexa-567-conjugated trastuzumab (Tz-568; 0.15 μg/ml) in cold Hank’s Balanced Salt Solution on ice for 1 h protected from light. Internalization was triggered with a temperature shift by adding pre-warmed serum-free media to the cells and incubating at +37°C for indicated times. 0 min sample serves as the no internalization control. The antibody internalization was stopped by washing with cold PBS and fixing with 4 % PFA for 10 min at RT. Fixed cells were washed and stored at +4°C in dark until imaging with 3i spinning disk confocal microscope.

### Immunohistochemistry, Ki-67 and TUNEL labelling

Formalin-fixed, paraffin-embedded tissue samples were cut to 4 µm sections, and deparaffinized and rehydrated with standard procedures. For immunohistochemistry of mouse xenografts, heat-mediated antigen retrieval was done for all samples in citrate buffer (pH 6). For Ki-67 labelling, sections were washed with washing buffer (Tris-HCl 0.05 M pH 7.6, 0.05 % Tween20) and Normal Antibody Diluent (NABD; Immunologic, BD09-125) and incubated with a Ki-67 antibody (14-5698, eBioscience, clone SolA15, diluted 1:2000) for 1 h. After washes, samples were incubated in 3 % H_2_O_2_ Tris-HCl for 10 min and washed again. Samples were incubated for 30 min with Rat probe (Biocare Medical rat on mouse HRP-polymer RT517 –kit), washed, and further incubated with R-O-M HRP-polymer (Biocare Medical rat on mouse HRP-polymer RT517 –kit) for 30 min. After washes, DAB solution (DAKO K3468) was added for 10 sec followed by washing. After counterstain with Mayer’s HTX, slides were dehydrated, cleared in xylene and mounted.

For TUNEL staining of apoptotic cells, endogenous peroxidase activity was blocked with incubation in 3% H2O2 in PBS for 15 min. The reaction mix containing recombinant terminal transferase (Cat No. 03 333 566 001, Roche), CoCl_2_ and Biotin-16-dUTP (11 093 070 910, Roche) in TdT buffer (1M Potassium cacodylate, 3% BSA in PBS, pH 6.6) was applied on slides and incubated for 1 h at +37ºC in a humidified chamber. As a positive control, one section was incubated for 30 min in DNAse solution at +37ºC before TUNEL staining, and as a negative control, the reaction mix was added without the transferase. Then the samples were incubated with End reaction-solution (300mM NaCl, 30 mM Sodium Citrate) for 30 min at RT and washed. Blocking was done with 3% BSA in PBS 45 min at RT after which the samples were incubated with ExtrAvidine (1:500 dilution in 1% BSA/PBS) for 30 min at +37ºC in a humidified chamber. After PBS washes, DAB solution (DAKO K3468) was added for 10 sec followed by washing. After counterstain with Mayer’s HTX, the slides were dehydrated, cleared in xylene, mounted and imaged with Pannoramic 250 Slide Scanner (3DHISTECH Ltd).

For bladder cancer tissue microarray (TMA) immunohistochemistry, BenchMark XT automated IHC/ISH slide staining system (Ventana Medical Systems, Inc.) with Cell Conditioning Solution (CC1) as a pretreatment was used for anti-HER2/neu (rabbit monoclonal, Ventana clone 4B5, 24 min incubation) and anti-EGFR (rabbit monoclonal, Ventana clone 5B7, 32 min), followed by UltraView Universal DAB Detection Kit (Ventana). For SORLA staining, antigen retrieval pretreatment was first done by microwaving the slides in citrate buffer (pH 6). Lab Vision autostainer (Thermo Fisher Scientific) with SORLA antibody (rabbit polyclonal, Atlas Antibodies; 1:300 dilution) was used and primary antibodies were detected with PowerVision Poly-HRP anti-mouse/anti-rabbit IHC system (Leica BioSystems). Finally, the slides were counterstained with hematoxylin.

IHC staining intensities of each tissue core were visually scored for SORLA as 0 (negative), 1 (weak to moderate), or 2 (strong), and for HER2 and EGFR as 0 (negative), 1 (weak), 2 (moderate) or 3 (strong). SORLA and HER2/EGFR stainings were evaluated independently by two readers and the mean and maximum values of three tissue cores were determined for each patient. In the final statistical analysis, maximum scores for all the stainings were used.

### Tissue microarray construction

Bladder cancer TMA construction was approved by the Research Ethics Board of the Hospital District of Southwest Finland (1.8.2006/301). FFPE tissue samples from consecutive 199 patients who underwent radical cystectomy in Turku University Hospital between 1985 and 2005 were used and three 1 mm tissue cores per patient were punched for TMA. Average age at cystectomy was 64 and none received neoadjuvant therapies.

### Quantification of intracellular HER2 levels

Several fields were randomly imaged with identical microscope settings. ImageJ was used for analysis and quantifications. Intracellular Tz-568 fluorescence signal was analysed from maximal intensity projections of 6 planes taken from the middle of the cell (determined by DAPI signal). Intracellular endogenous HER2 fluorescence intensity was measured from a single middle plane of the cell. Intracellular signal was quantified by manually gating the intracellular part of the cell and the intracellular signal was measured and normalized to the cell area. Results are pooled from two independent biological replicates.

### Quantification of LAMP1 positive late-endosome/lysosome aggregation

SORLA silenced and control silenced cells were fixed and stained for LAMP1 (Santa-Cruz, SC-20011) and DAPI as described above. Cells were imaged with identical microscope settings with Carl Zeiss LSM780 laser scanning confocal microscope. The image processing and quantifications were performed with Image J software. The level of LAMP1 positive late-endosome/lysosome aggregation was quantified from the middle plane. To distinguish between individual late-endosome/lysosome aggregates, watershed segmentation was used before quantification. The areas of late-endosome/lysosome aggregates were quantified from a single cell. LAMP1 positive area larger than 5 µm^2^ was considered as aggregate. Total area of LAMP1 positive late-endosomes/lysosomes was calculated based on LAMP1 staining as well as the area of aggregates larger than 5 µm^2^. The percentage of lysosomal aggregation was determined by the ratio of area of aggregates larger than 5 µm^2^ to total LAMP1 positive late-endosome/lysosome area.

### Quantification of Ki-67 and TUNEL positivity in 5637 cell xenografts

Pannoramic viewer (3DHISTECH Ltd) was used to scan histology slides and export images (2x magnification) for image processing and quantifications, which were performed identically for all the samples with ImageJ software. The TUNEL and Ki-67 positive tumor areas were thresholded with MaxEntropy and quantified. The ratio of Ki-67 positive (or TUNEL positive) area to total tumor area was calculated.

### Co-localization analysis

Pixel-intensity based Pearson correlation coefficient (R) between two channels was calculated using *coloc2* plugin (https://imagej.net/Coloc_2) of ImageJ v1.51s with default parameters. Percentage colocalization between the vesicular particles was done using ComDet 0.3.6.1 (https://github.com/ekatrukha/ComDet) plugin of ImageJ v1.51s. Particles were detected in both channels independently at approximated particle sizes of 4 pixels with sensitivities of signal/noise ratio of 4. Colocalization was determined based on a maximum distance between two particle centers of 5 pixels and expressed as percentage.

### Statistical analysis

The GraphPad Prism software and two-tailed Student’s *t*-test (paired or unpaired, as appropriate) was used for statistical analysis. Normal distribution of the data was tested with Shapiro-Wilk normality test. When data was not normally distributed, a Mann-Whitney test was used. In proliferation assays, two-way ANOVA was used. P-values <0.1 are shown in graphs.

